# Polyphyletic ancestry of expanding Patagonian Chinook salmon populations

**DOI:** 10.1101/079459

**Authors:** Cristian Correa, Paul Moran

## Abstract

Chinook salmon native to North America are spreading through South America’s Patagonia and have become the most widespread anadromous salmon invasion ever documented. To better understand the colonization history and role that genetic diversity might have played in the founding and radiation of these new populations, we characterized ancestry and genetic diversity across latitude (39-48°S). Samples from four distant basins in Chile were genotyped for 13 microsatellite loci, and allocated, through probabilistic mixture models, to 148 potential donor populations in North America representing 46 distinct genetic lineages. Patagonian Chinook salmon clearly had a diverse and heterogeneous ancestry. Lineages from the Lower Columbia River were introduced for salmon open-ocean ranching in the late 1970s and 1980s, and were prevalent south of 43°S. In the north, however, a diverse assembly of lineages was found, associated with net-pen aquaculture during the 1990s. Finally, we showed that possible lineage admixture in the introduced range can confound allocations inferred from mixture models, a caveat previously overlooked in studies of this kind. While we documented high genetic and lineage diversity in expanding Patagonian populations, the degree to which diversity drives adaptive potential remains unclear. Our new understanding of diversity across latitude will guide future research.

## Introduction

Multiple independent introduction events can lead to shifts in genetic variation relative to native source populations, potentially boosting invasiveness and potential for rapid local adaptation ^1–4^. Furthermore, different introduction vectors delivering distinct genetic lineages in different regions can result in a mosaic of populations with varying genetic diversity and evolutionary potential ^1,2,5,6^. We investigated self-sustaining Chinook salmon [*Oncorhynchus tshawytscha* (Walbaum)] populations, currently part of a rapid colonization that is sweeping through Patagonia, the binational region of Chile and Argentina at the southern cone of South America. Specifically, we measured genetic diversity and evaluated the most likely phylogenetic origins of four self-sustaining populations spread over a wide geographical range that received multiple distinct introductions from different kinds of artificial propagation.

Beginning in the 1870s, Chinook salmon, native to North America and the North Pacific Ocean, were deliberately introduced into innumerable rivers in all continents except Antarctica ^7^; yet successful naturalization has been rare. Self-sustaining adfluvial (migrating between lake and river) Chinook salmon populations have been established in the North American Great Lakes ^8^, but anadromous (migrating between the sea and river) populations outside their native range exist only in New Zealand's South Island and in South America's Patagonia ^7^. The phylogenetic ancestry of New Zealand Chinook salmon was tracked to introductions in the early 1900s from the Sacramento River fall run (seasons characterize typical adult return to freshwater), most likely Battle Creek, California ^9–11^. Originally stocked in one river of the South Island, the fish naturalized and within a decade expanded their range considerably ^11–13^. When studied about 30 salmon generations later, a number of phenotypic traits had evolved apparently in response to local environmental conditions ^11,14^, offering increased local fitness ^15^. The monophyletic ancestry and known history of introduction in New Zealand greatly facilitated that research ^11,15^. This rapid evolution in New Zealand underscores the potential of Chinook salmon to adapt to new environments, yet it begs the question of why successful introductions have been rare elsewhere.

Patagonian Chinook salmon are much less studied than New Zealand Chinook salmon. Patagonian populations are younger than those in New Zealand but much more widespread. Self-sustaining populations occur in a vast region on western slopes south of 38°S draining through Chile and into the Pacific Ocean. They are also established on eastern slopes south of 50°S draining through Argentina and into the Atlantic Ocean (Figure 1). The ancestral origin of these populations is unclear due to fragmentary historical records and insufficient study, and numerous potential donor populations must be considered ^16–19^.

**Figure 1:**
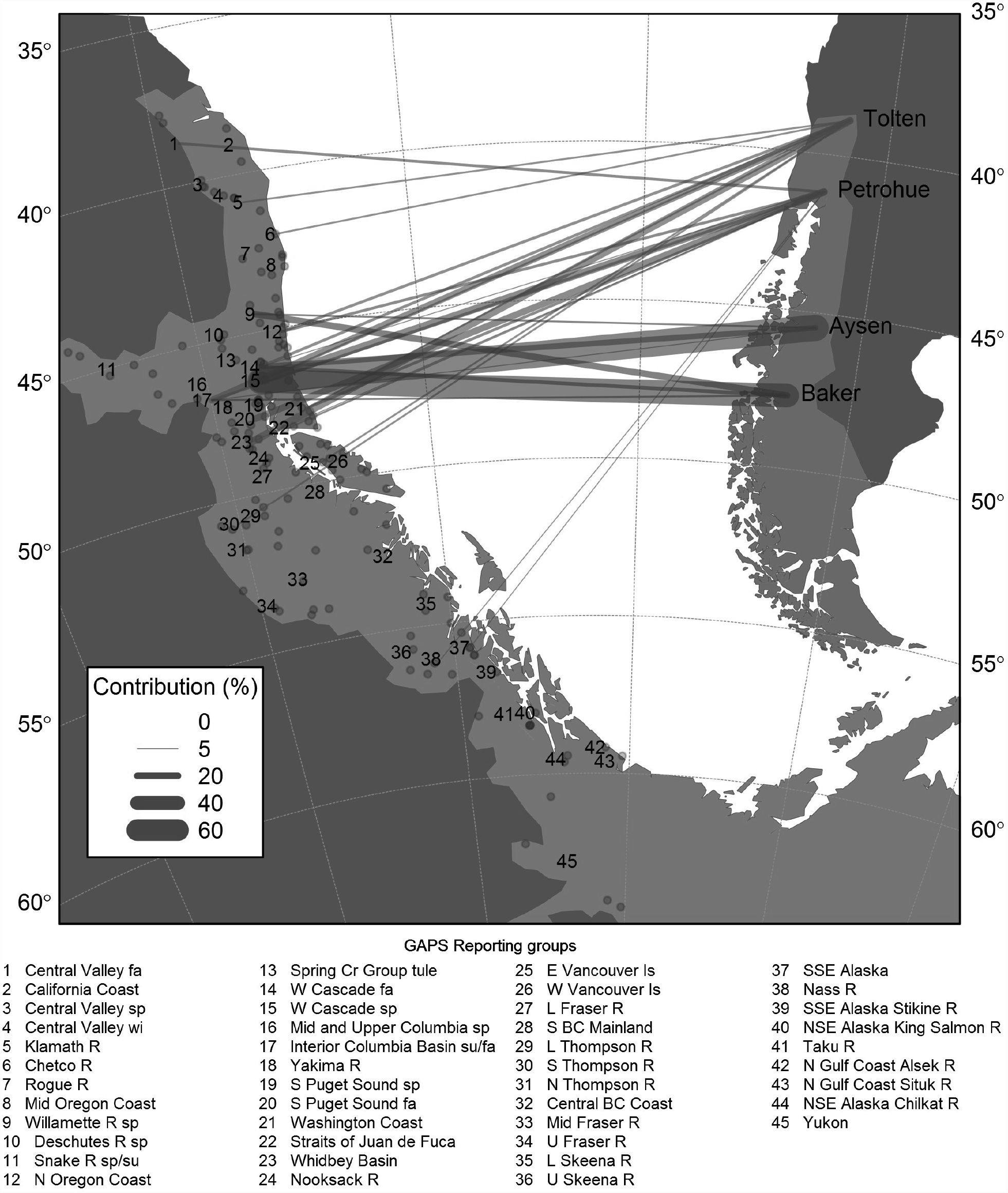
Chinook salmon native (green) and invasive (brown) range in the Americas, and illustration of genetic contribution of North American lineages to South American introduced populations (connecting vectors). The North American continent has been rotated and translated such that the North American and South American Pacific coasts face each other matching latitudes (longitudes were shifted for clarity). Connecting vectors represent average genetic contribution (proportional to line thickness) of the 46 North American baseline reporting groups (lineages) used in conditional maximum likelihood (CML) mixture analysis. Reporting groups were characterized by 146 baseline populations (dots).

Correa and Gross ^18^ reviewed the historical records of introductions into Latin America, records of salmon in the wild, locations of fish farming, and their own field observations. They concluded that, out of the many introduction attempts beginning in the late 1800s, two relatively recent experimental commercial open-ocean ranching operations were most likely the sources of the introduced populations. For the purpose of this study, we distinguish two classes of artificial propagation. In open-ocean ranching, returning adults are spawned artificially in fish hatcheries and the juveniles are released to grow at sea. These free-ranging fish may be harvested at sea but especially in a terminal fishery of maturing adults homing back to the stream of release. The second class is floating net-pen aquaculture. In this case, fish are held captive from hatching until harvest or reproduction (see below). One open-ocean ranching operation took place on Chiloé Island (42°S) during the period 1978-1982 using primarily Cowlitz River Hatchery spring-run brood stock from the Lower Columbia River, Washington, U.S.A. ^20^. The other open-ocean ranching operation took place further south in the Magellan region (51°S) during the period 1982-1989 using University of Washington Hatchery fall-run broodstock, Puget Sound, Washington, as well as the progeny of adult fish returning to Chiloé Island ^21–23^.

By 1988, Chilean salmonid propagation shifted entirely from open-ocean ranching to net-pen aquaculture in captivity. Despite the captive design, fish escaped in the thousands every year and many of those escapes were poorly documented (Soto et al. 2001; Naylor et al. 2005; Buschmann et al. 2006; Arismendi et al. 2009). The stocks utilized, also poorly documented, were relatively diverse, as ova importations came at least from Washington, Oregon, Vancouver Island, and from New Zealand (the latter with ancestry from the Sacramento River, Central Valley, California) ^24^. In contrast to open-ocean ranching, net-pen aquaculture was mostly concentrated in Chile's Lake District region (42°S). A summary of Chinook salmon introductions to Patagonia is presented in Table 1. A complete review of introductions is provided as an online supplement (Supplementary Table S1).

**Table 1:**
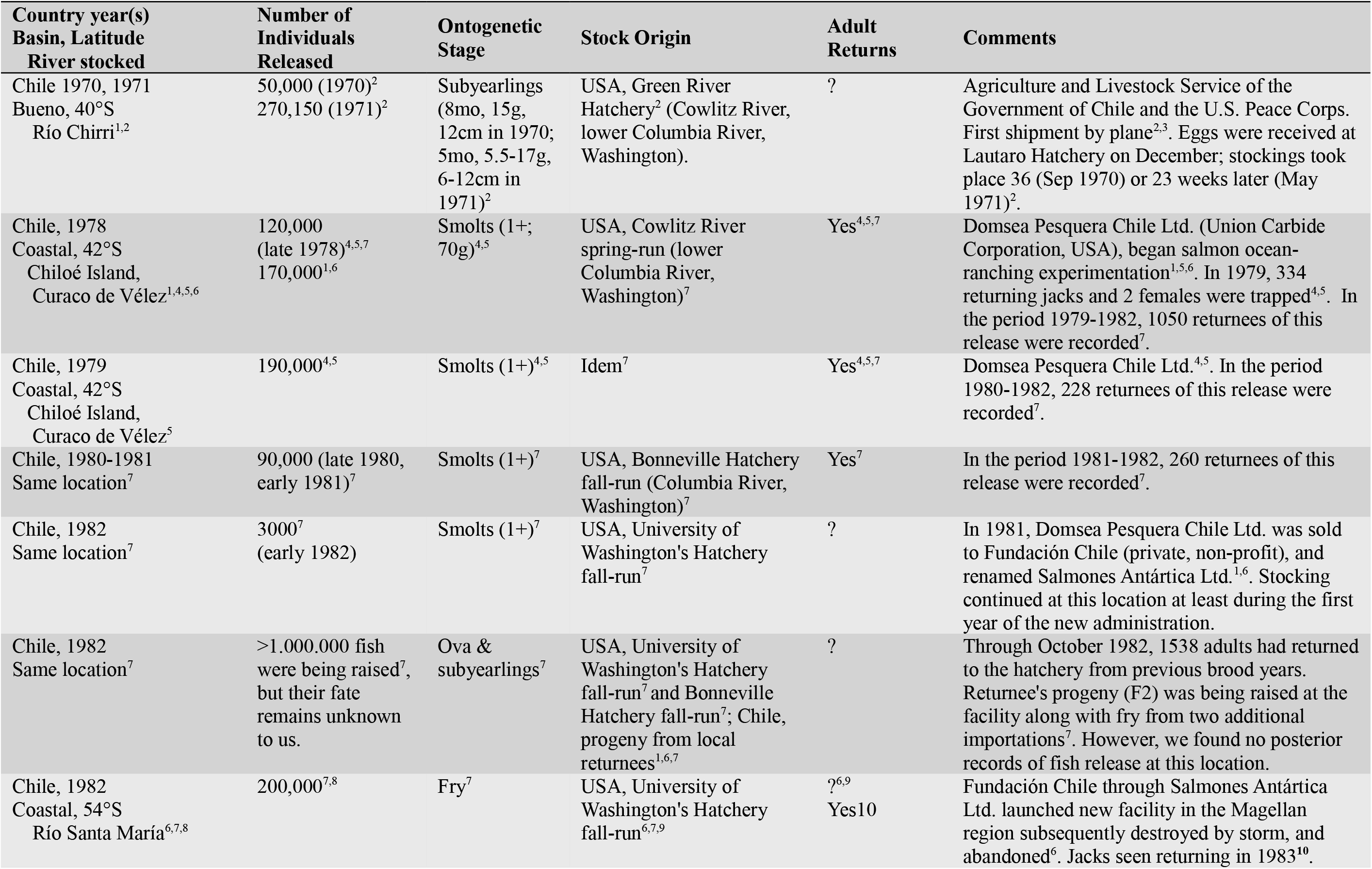
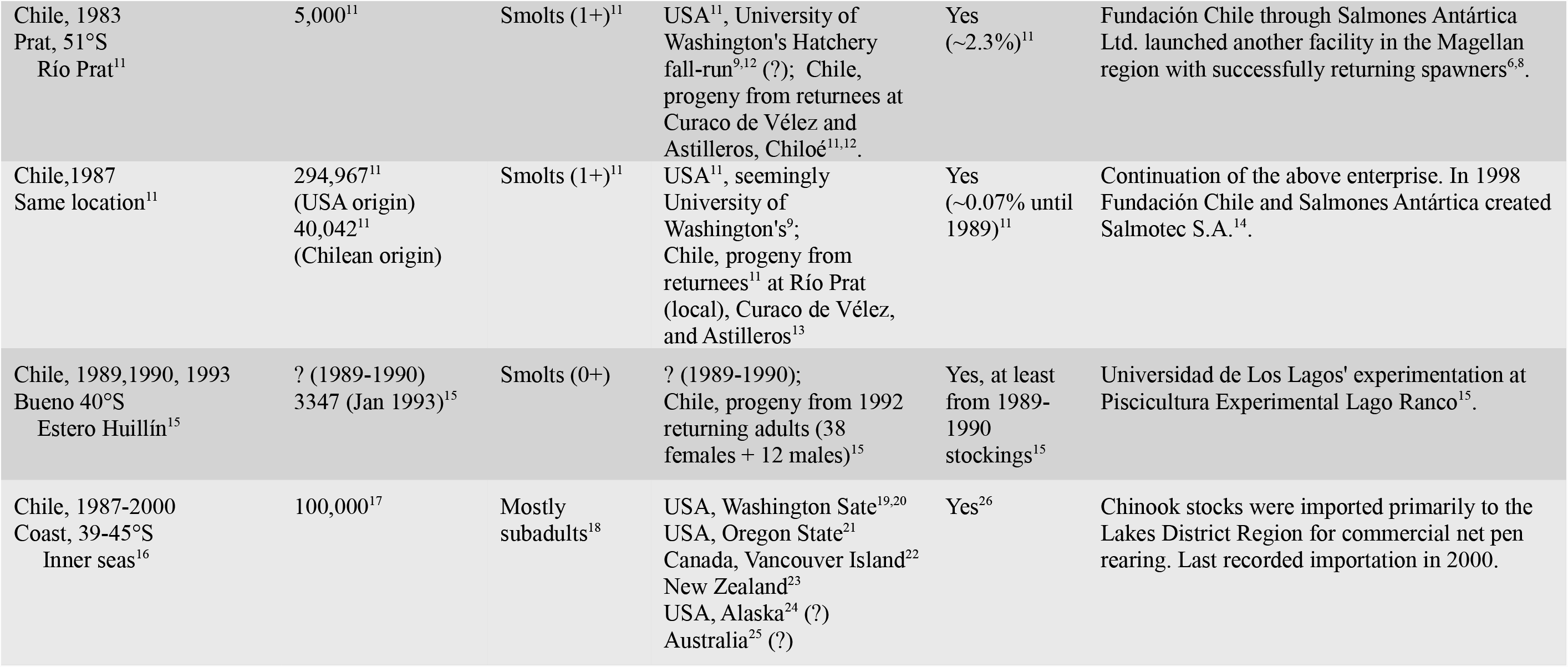
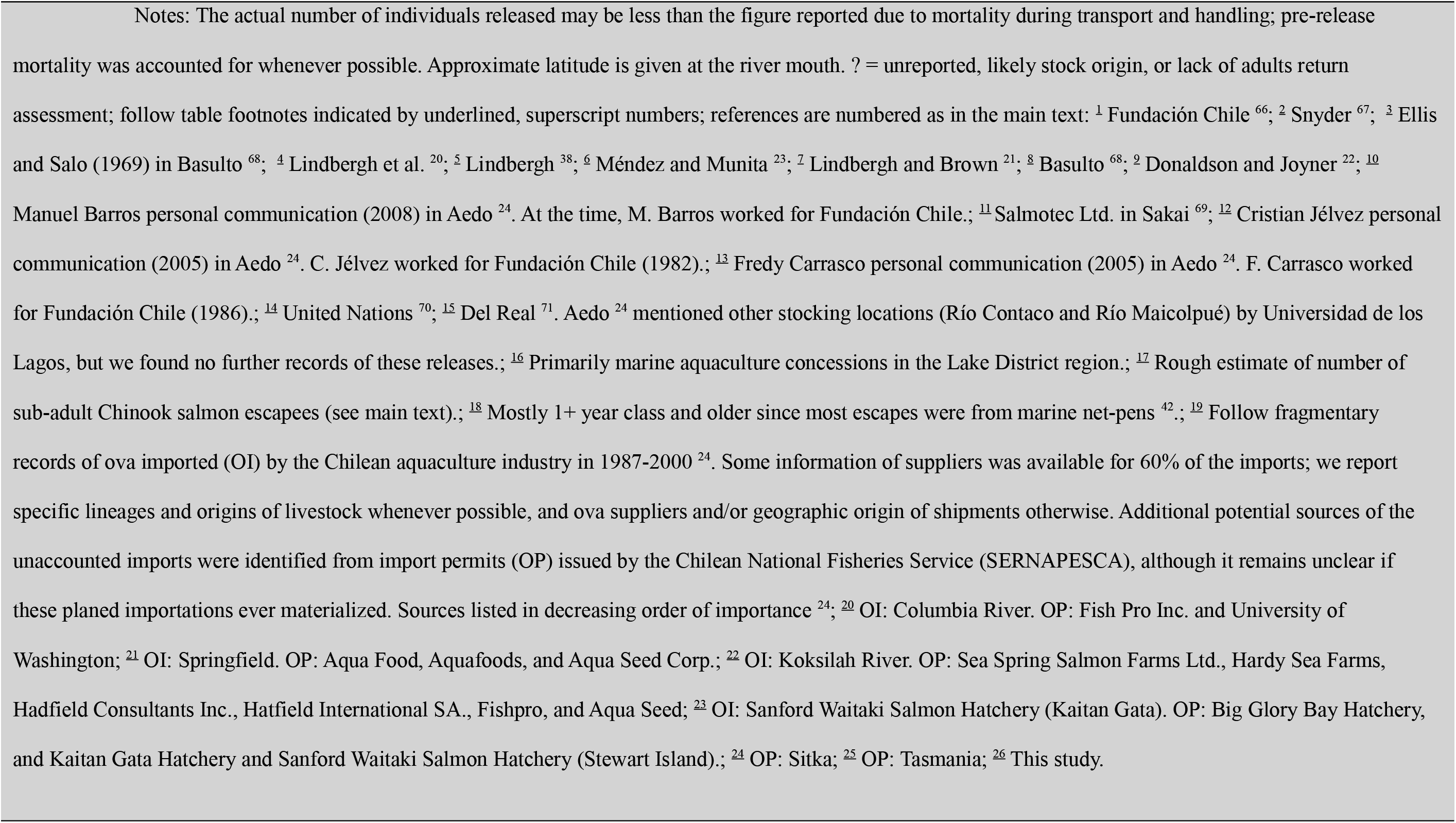
Chinook salmon deliberate and accidental releases in Patagonia since 1970 (a complete review spanning earlier introductions is available in Supporting Information, Table S1; modified from Correa and Gross 2008).

| Country year(s)<br>Basin, Latitude<br>River stocked | Number of<br>Individuals<br>Released | Ontogenetic<br>Stage | Stock Origin | Adult<br>Returns | Comments |
| --- | --- | --- | --- | --- | --- |
| Chile 1970, 1971<br>Bueno, 40°S<br>Río Chirri<sup>1,2</sup> | 50,000 (1970)<sup>2</sup><br>270,150 (1971)<sup>2</sup> | Subyearlings<br>(8mo, 15g,<br>12cm in 1970;<br>5mo, 5.5-17g,<br>6-12cm in<br>1971)<sup>2</sup> | USA, Green River<br>Hatchery<sup>2</sup> (Cowlitz River,<br>lower Columbia River,<br>Washington). | ? | Agriculture and Livestock Service of the<br>Government of Chile and the U.S. Peace Corps.<br>First shipment by plane<sup>2,3</sup>. Eggs were received at<br>Lautaro Hatchery on December; stockings took<br>place 36 (Sep 1970) or 23 weeks later (May<br>1971)<sup>2</sup>. |
| Chile, 1978<br>Coastal, 42°S<br>Chiloé Island,<br>Curaco de Vélez<sup>1,4,5,6</sup> | 120,000<br>(late 1978)<sup>4,5,7</sup><br>170,000<sup>1,6</sup> | Smolts (1+;<br>70g)<sup>4,5</sup> | USA, Cowlitz River<br>spring-run (lower<br>Columbia River,<br>Washington)<sup>7</sup> | Yes<sup>4,5,7</sup> | Domsea Pesquera Chile Ltd. (Union Carbide<br>Corporation, USA), began salmon ocean-<br>ranching experimentation<sup>1,5,6</sup>. In 1979, 334<br>returning jacks and 2 females were trapped<sup>4,5</sup>. In<br>the period 1979-1982, 1050 returnees of this<br>release were recorded<sup>7</sup>. |
| Chile, 1979<br>Coastal, 42°S<br>Chiloé Island,<br>Curaco de Vélez<sup>5</sup> | 190,000<sup>4,5</sup> | Smolts (1+)<sup>4,5</sup> | Idem<sup>7</sup> | Yes<sup>4,5,7</sup> | Domsea Pesquera Chile Ltd.<sup>4,5</sup>. In the period<br>1980-1982, 228 returnees of this release were<br>recorded<sup>7</sup>. |
| Chile, 1980-1981<br>Same location<sup>7</sup> | 90,000 (late 1980,<br>early 1981)<sup>7</sup> | Smolts (1+)<sup>7</sup> | USA, Bonneville Hatchery<br>fall-run (Columbia River,<br>Washington)<sup>7</sup> | Yes<sup>7</sup> | In the period 1981-1982, 260 returnees of this<br>release were recorded<sup>7</sup>. |
| Chile, 1982<br>Same location<sup>7</sup> | 3000<sup>7</sup><br>(early 1982) | Smolts (1+)<sup>7</sup> | USA, University of<br>Washington's Hatchery<br>fall-run<sup>7</sup> | ? | In 1981, Domsea Pesquera Chile Ltd. was sold<br>to Fundación Chile (private, non-profit), and<br>renamed Salmones Antártica Ltd.<sup>1,6</sup>. Stocking<br>continued at this location at least during the first<br>year of the new administration. |
| Chile, 1982<br>Same location<sup>7</sup> | >1.000.000 fish<br>were being raised<sup>7</sup>,<br>but their fate<br>remains unknown<br>to us. | Ova &<br>subyearlings<sup>7</sup> | USA, University of<br>Washington's Hatchery<br>fall-run<sup>7</sup> and Bonneville<br>Hatchery fall-run<sup>7</sup>; Chile,<br>progeny from local<br>returnees<sup>1,6,7</sup> | ? | Through October 1982, 1538 adults had returned<br>to the hatchery from previous brood years.<br>Returnee's progeny (F2) was being raised at the<br>facility along with fry from two additional<br>importations<sup>7</sup>. However, we found no posterior<br>records of fish release at this location. |
| Chile, 1982<br>Coastal, 54°S<br>Río Santa María<sup>6,7,8</sup> | 200,000<sup>7,8</sup> | Fry<sup>7</sup> | USA, University of<br>Washington's Hatchery<br>fall-run<sup>6,7,9</sup> | ?<sup>6,9</sup><br>Yes<sup>10</sup> | Fundación Chile through Salmones Antártica<br>Ltd. launched new facility in the Magellan<br>region subsequently destroyed by storm, and<br>abandoned<sup>6</sup>. Jacks seen returning in 1983<sup>10</sup>. |
| Chile, 1983<br>Prat, 51°S<br>Río Prat<sup>11</sup> | 5,000<sup>11</sup> | Smolts (1+)<sup>11</sup> | USA<sup>11</sup>, University of Washington's Hatchery fall-run<sup>9,12</sup> (?); Chile, progeny from returnees at Curaco de Vélez and Astilleros, Chiloé<sup>11,12</sup>. | Yes<br>(~2.3%)<sup>11</sup> | Fundación Chile through Salmones Antártica Ltd. launched another facility in the Magellan region with successfully returning spawners<sup>6,8</sup>. |
| Chile, 1987<br>Same location<sup>11</sup> | 294,967<sup>11</sup><br>(USA origin)<br>40,042<sup>11</sup><br>(Chilean origin) | Smolts (1+)<sup>11</sup> | USA<sup>11</sup>, seemingly University of Washington's<sup>9</sup>; Chile, progeny from returnees<sup>11</sup> at Río Prat (local), Curaco de Vélez, and Astilleros<sup>13</sup> | Yes<br>(~0.07% until 1989)<sup>11</sup> | Continuation of the above enterprise. In 1998 Fundación Chile and Salmones Antártica created Salmotec S.A.<sup>14</sup>. |
| Chile, 1989, 1990, 1993<br>Bueno 40°S<br>Estero Huillín<sup>15</sup> | ? (1989-1990)<br>3347 (Jan 1993)<sup>15</sup> | Smolts (0+) | ? (1989-1990); Chile, progeny from 1992 returning adults (38 females + 12 males)<sup>15</sup> | Yes, at least from 1989-1990 stockings<sup>15</sup> | Universidad de Los Lagos' experimentation at Piscicultura Experimental Lago Ranco<sup>15</sup>. |
| Chile, 1987-2000<br>Coast, 39-45°S<br>Inner seas<sup>16</sup> | 100,000<sup>17</sup> | Mostly subadults<sup>18</sup> | USA, Washington State<sup>19,20</sup><br>USA, Oregon State<sup>21</sup><br>Canada, Vancouver Island<sup>22</sup><br>New Zealand<sup>23</sup><br>USA, Alaska<sup>24</sup> (?)<br>Australia<sup>25</sup> (?) | Yes<sup>26</sup> | Chinook stocks were imported primarily to the Lakes District Region for commercial net pen rearing. Last recorded importation in 2000. |

Therefore, in contrast to New Zealand, Patagonian Chinook salmon represent a potentially polyphyletic assemblage, offering new and interesting research avenues to study colonization dynamics and contemporary evolution. Although our results are mostly concordant with the few available molecular studies regarding polyphyletic ancestry, we identified a somewhat different array of contributing lineages. We also identified a methodological anomaly not previously recognized in studies of this kind.

We analysed the genetic diversity and population structure of Chinook salmon in four western Patagonia watersheds. We then compared these to native North America populations, and finally estimated ancestry and lineage distribution in regions that differed in their history of artificial propagation by open-ocean ranching and net-pen aquaculture. For estimation of ancestry, we used two different classes of genetic mixture analysis, conditional maximum likelihood (CML) and model-based clustering (M-BC). We took advantage of a large, interagency, microsatellite baseline as our known-origin reference against which we compared fish from Patagonia. The international Genetic Analysis of Pacific Salmonids (GAPS) consortium assembled a comprehensive genetic baseline for coast-wide fishery management of mixed-stock fisheries ^25^. The accuracy and precision of mixture analysis using the GAPS baseline is substantial with either conditional or unconditional Bayesian methods ^25–28^, and provides a potentially useful approach for inferring the ancestry of Patagonian Chinook salmon. CML mixture analysis was used in two recent studies of the ancestry of Patagonian Chinook salmon ^29,30^. However, as far as we know, the validity of this application in mixed-origin populations has not been demonstrated. Introduced populations violate the fundamental assumption of the model—that individuals in the unknown “mixture” actually originated from one of the baseline populations. Newly founded populations are isolated and expected to diverge from their ancestors through founder effects and genetic drift ^29^. An even greater concern is interbreeding of mixed-origin fish in the new range (admixture), producing novel genotypes not accurately attributable to any single source population. Founding effects and drift are not easily accounted for, but neither are they likely to fundamentally confound our analysis. When considering microsatellites or other neutral markers, it is unlikely that founding and drift would result in a Patagonian population looking more like an unrelated population than the true population of origin. Such a result would require parallel patterns of allele frequency convergence at multiple highly polymorphic loci. The problem of interbreeding is more serious. It was not clear to us how the Rannala and Mountain ^31^ CML mixture algorithm would treat individuals of mixed origin. Simulations and sensitivity analysis helped us answer that question, and M-BC allowed an independent evaluation of CML mixture analysis in this unusual application of stock identification of introduced and naturalized populations.

## Results

### Data quality, neutrality, and genetic structure

Data quality and genotyping success were high, with an average of 12 out of 13 microsatellite loci scored per individual fish. We removed from the analysis 9 Toltén individuals that gave no reliable genotypes (collected as decomposing carcass samples), leaving 87 individual Patagonian Chinook salmon for our study.

*Ots*213 departed significantly from neutral expectation (*F*_ST_ outlier test) and was removed from further population genetic analyses due to potential directional selection (but was retained for genetic mixture analysis, which is generally robust to departures from neutrality). *Oki*100 and *Ots*208b also departed from neutral expectation but were only marginally significant and so were retained. None of these loci are known to diverge from neutral expectation in North American populations ^32^. Pairwise allele frequency differences were not significant between sample collection locations within river basins in Patagonia (range of intrabasin P values, 0.0042 – 0.690, Bonferroni-corrected α = 0.0014) and were therefore pooled for population-level analyses (but see below). The analyses we present here are based on those river-basin-level aggregates of collections that are intended to represent separate populations, though because our samples are small, we tested and evaluated that assumption in several ways. For example, mean *F*_IS_ values for Patagonian populations (0.013, SD = 0.0195) were substantially larger than estimates from the North America baseline populations (0.008, SD = 0.0159, Supplementary Table S2). Those heterozygote deficits in Patagonia might indicate a Wahlund effect of having sampled distinct populations within a river basin. Although departures from Hardy-Weinberg expectations were non-significant, we recognize there was little power with such small sample size. Non-significant heterozygote deficits were observed at *Ots*201 and *Ots*213 (again, *Ots*213 was not included in most population genetic analyses because of its highly significant departure from neutral expectation).

### Genetic diversity

Average genetic diversity in the Patagonian populations (mean *H*_S_ = 0.83, SD = 0.024) was essentially identical to to the average of all North American baseline populations (0.83, SD = 0.038) and close to the populations identified as likely founders (see below, *e*.g., Puget Sound, Whidbey Basin, West Cascades, and Oregon Coast, 0.86, SD = 0.026, Supplementary Table S2). Similarly, average allelic richness in Patagonian populations (*AR* = 9.3, SD = 1.48) was not significantly different from that observed in all North American populations (9.8, SD = 1.13) nor in the putative founders (10.6, SD = 0.70). The 259 alleles observed in Patagonia included more than half of the 490 observed in North America, from the Yukon River in the Bering Sea to Central Valley California—most of the native species range ^32^. Within Patagonia, overall genetic diversity did not differ significantly among river basins, as inferred from heterozygosity and allelic richness (Supplementary Table S2). Even though our population samples from Patagonia were small, we seem to have been successful in capturing a broad cross section of genetic diversity in these introduced populations.

### Population structure

Between Patagonian river basins, allele frequencies were highly significantly different (mean pair-wise P = 0.00017, where Bonferroni-corrected α = 0.00833). Similarly, pairwise *F*_ST_ values were large among Patagonian populations (mean *F*_ST_ = 0.039, SD = 0.004), especially considering the relatively narrow range of genetic differentiation among the putative source populations (0.028, SD = 0.006) relative to all of North America (0.063, SD = 0.01).

Contingency testing, pair-wise *F*_ST_ estimates, and jack knife self-assignment analysis were all consistent in showing the closest genetic relationship between the two southern populations Aysén and Baker (*F*_ST_ = 0.01). Petrohué was similar to Baker and Aysén, whereas Toltén, the northern-most population, was the most distinct of the four [i.e., in Newick format: ((Aysén, Baker) Petrohué) Toltén].

### Origin of Patagonian Chinook salmon (CML mixture analysis)

We found evidence of multiple North American lineages in Patagonia. Of the 46 reporting groups, 16 (35%) were identified by CML mixture analysis as possibly represented in Patagonia (Figure 1 and Figure 2). However, based on our simulations, we expected a small fraction of spurious assignments/allocation associated with admixture among lineages occurring in the introduced range (see below, *Results: Simulated mixed-origin founding*). Hence, in order to emphasize major contributing lineages and avoid over-interpretation, we focused on putative contributors with *c.* 10% or greater allocation from any source to any single Patagonian population (Figure 2; see below *Simulated mixed-origin founding*). Seven potential source lineages satisfied this criterion. Approximately south to north in their native range, these were: Willamette River spring, North Oregon Coast, West Cascade fall, West Cascade spring, Interior Columbia Basin summer/fall, South Puget Sound fall, and Whidbey Basin.

**Figure 2:**
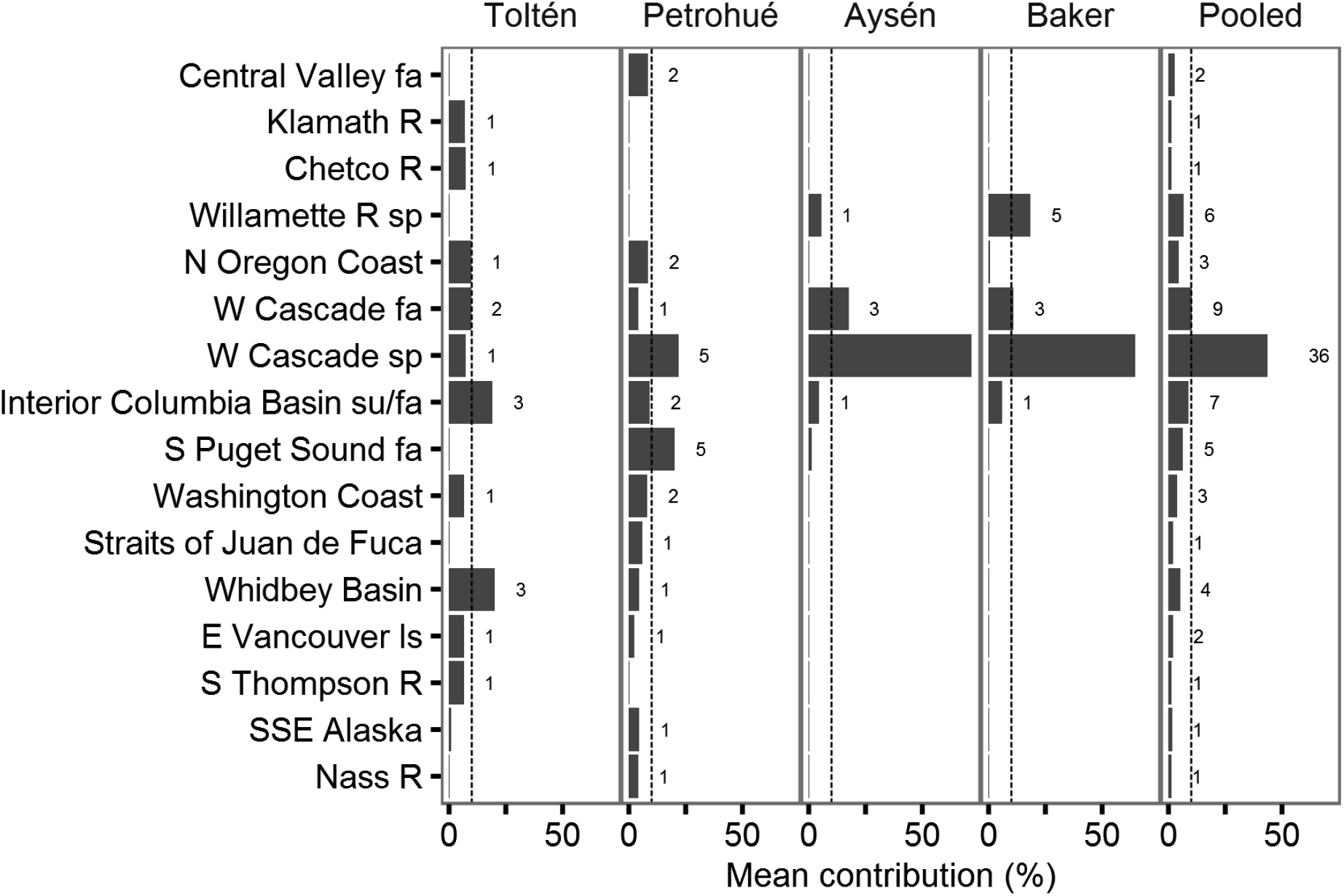
Estimated contribution of North American Chinook salmon lineages to Patagonian populations, as inferred from reporting group-level CML. A cline in lineage diversity was apparent, with populations from southern rivers (Aysén and Baker) dominated by the West Cascade spring run lineage, whereas those from northern rivers (Toltén and Petrohué) a broad diversity of lineages. With a few exceptions, contributing lineages were concordant with historical introductions and regional differences in propagation activities. Numbers beside bars of mean proportional genetic contribution indicate number of individuals allocated to a particular lineage. Dashed vertical line indicate 10% genetic contribution.

The estimated number of lineages that contributed to our South American populations was higher in the north (Toltén and Petrohué) than in the south of our study area (Aysén and Baker) (Figure 1). West Cascade fall, and especially West Cascade spring, contributed substantially, especially in the south with 71% and 65% genetic contribution of West Cascade spring to Aysén and Baker populations (Figure 2). Interior Columbia basin summer/fall lineage contributed to all four Patagonian populations, but especially to those in the north. Other contributors showed more localized effects. For example, South Puget Sound fall in Petrohué (20%), Whidbey Basin in Toltén (20%), and Willamette River spring in Baker (18%). Various other donors had lower contributions, but, again, some of those results might be attributable to the misassignments we observed in the simulation of mixed ancestry.

Whereas the two northern river basins (39°-42°S) appeared highly polyphyletic, the two southern basins (45°-48°S) were nearly monophyletic, attributing nearly all of their ancestry to the closely related lower Columbia River, West Cascade spring and Willamette River spring lineages. Despite small sample sizes and apparent mixed ancestry, we found general agreement between modeled stock composition estimates and the proportions of individual fish assignments (Figure 2; Supplementary Table S3).

### Split assignments consistent with primary donors

The ancestries of most fish were readily identifiable though CML mixture analysis. Most assigned with high probability to one of the baseline reporting groups (average maximum *a posteriori* value = 0.903, SD = 0.145) or reference populations (0.861, SD = 0.168) (Supplementary Figure S1). High assignment probabilities to baseline reporting groups were observed even in watersheds where spawners from multiple lineages were collected together (*i.e.*, Toltén and Petrohué). Both at the reporting group and population-level, individual fish that did show affinity to multiple source lineages, were invariably associated with the same sources to which other fish in the same collection assigned with high probability. For example, Petrohué and Toltén had individuals that assigned with relatively high probability to North Oregon Coast, West Cascade spring and fall, Interior Columbia Basin summer/fall, and South Puget Sound fall; however other presumed mixed-origin individuals split their assignment probability among those same sources.

### Simulated mixed-origin founding

Our simulation result showed that most individual fish in a population derived from multiple sources would assign back to those sources, often splitting their assignment probability between source populations (Supplementary Figure S2). However, we also learned that sometimes a non-trivial number of simulated, mixed-origin fish might assign with high probability to unrelated lineages. Some misassignment was expected to genetically similar reporting groups. Both reporting groups used in our simulation have closely related sister groups in the GAPS baseline. West Cascade spring is genetically similar to West Cascade fall, and South Puget Sound fall is similar to Whidbey Basin (an adjoining inland water body). Most simulated fish (61.3%) assigned to one or the other of the true source reporting groups. Another 28.2% assigned to those closely related sister groups. That level of misassignment was expected based on leave-one-out jackknife analysis of the North American baseline. Approximately 17% of real fish collected from Cowlitz River spring (one of the seed populations for the simulation) misassigned to Cowlitz River fall in the West Cascade fall reporting group.

Expected misassignment to closely related reporting groups contrasted strongly with assignment of simulated individuals to unrelated groups. In our simulation, 10% of individuals assigned to unrelated reporting groups, 44% of these with high assignment probability (*P* ≥ 0.8). Interior Columbia Basin summer/fall received about half of these misassignments, whereas the rest were distributed among 14 other unrelated reporting groups. This level of misassignment observed in simulated fish was much higher than the normal level of misassignment observed between closely related reporting groups in North America. Note that this result is completely simulated and is not related to mutation or drift in the founded populations or in the baseline.

Our simulation study confirmed the general utility of CML mixture analysis for individual assignment and proportional allocation despite admixture, but also showed the potential for misleading results, even when individuals assign with high probability. Therefore, we interpreted our empirical results of CML mixture analysis of real fish with caution when the estimated proportional contribution of a reporting group approached the level of misassignment we observed in the simulation (i.e., <10%, see above *Results: Origin of Patagonian Chinook salmon*). Any criterion would be somewhat arbitrary, but for the empirical dataset there seemed to be a break between 0.08 and 0.1, and we knew that values below 0.08 could be relatively strongly influenced by the spurious assignments we revealed in the simulation.

Therefore, the simulations largely confirmed the utility of the CML algorithm for studies of mixed ancestry with the important caveat that a few mixed origin individuals can show high probability of membership to completely unrelated lineages.

### Equivocal results regarding expected contributors

Not a single fish we sampled from four locations in Patagonia showed any affinity whatever to the University of Washington Hatchery fall-run stock (zero relative probability of membership). Only five fish, collected in the Petrohué basin, assigned with high probability to another Puget Sound fall-run hatchery stock, Soos Creek (from which University of Washington broodstock was derived), but none were similar to our contemporary sample from University of Washington Hatchery fall-run stock.

Central Valley California was also a suspected source of Patagonian Chinook salmon, however, our results offered very little support for that conclusion at the locations we sampled. Only two fish assigned to Central Valley populations, one to Tuolumne River and the other to Stanislaus River. It was unclear if those two fish demonstrated true Central Valley ancestry or spurious assignments resulting from admixture (see above *Simulated mixed-origin founding*).

### Model-based clustering (M-BC)

Our reduced reference dataset for M-BC (see *Methods*) included 31 of the GAPS baseline populations ^32^ from seven reporting groups (Whidbey Basin, S. Puget Sound fall, Interior Columbia Basin summer/fall, West Cascade fall, West Cascade spring, North Oregon Coast, and Willamette River spring). These reporting groups contributed most (84.13%) of the combined posterior probability in the reporting group-level CML mixture analysis (Figure 2). Examination of log likelihood of the modelled clusters ^33^ for various numbers of clusters (*K*) in the North American populations showed a peak in delta log likelihood at *K* = 3 and *K* = 5 (Supplementary Figure S3). We observed the clearest discrimination of the seven reporting groups at *K* = 5 (Supplementary Figure S4), and that value showed high concordance with the genetic stock reporting groups that we used in the CML mixture analysis (Figure 3a). Concordance between M-BC and CML was also evident in the Patagonian populations where the mean *Q*-values matched the proportional estimates of North American lineages based on CML (compare Figure 2 with Figure 3). For example, from south to north, individual Baker fish showed *Q*-value proportions mostly consistent with ancestry from West Cascade spring/fall (Q3), or a combination of West Cascade spring and Willamette River spring runs (Q4). Aysén fish mostly showed affinity with West Cascade spring/fall runs (Q3), but also some possible Willamette River spring (Q4) and little South Puget Sound fall and Whidbey Basin influence (Q5). In Petrohué, West Cascade spring/fall and Willamette River (Q3 and Q4) lineages were still important, but a substantial increase of all other lineages was apparent, particularly South Puget Sound fall and Whidbey Basin (Q5). This trend towards more diverse ancestry in the north of the Patagonian range persisted in Toltén where fish showed affinity to all five ancestral lineages inferred by the M-BC analysis (Q1 to Q5), and by extension the seven lineages distinguished in phylogeographic analysis of Chinook salmon in the native range ^32^. Thus, we observed a spatial gradient in the ancestry of Patagonian Chinook salmon populations (Figure 3b). In general, model based clustering suggested broader, more uniform contributions from the source populations that were identified with CML mixture analysis. Beyond that, however, results from M-BC were highly concordant with CML and led to nearly identical inferences about the ancestry of naturally-sustaining, Patagonian Chinook salmon populations.

**Figure 3:**
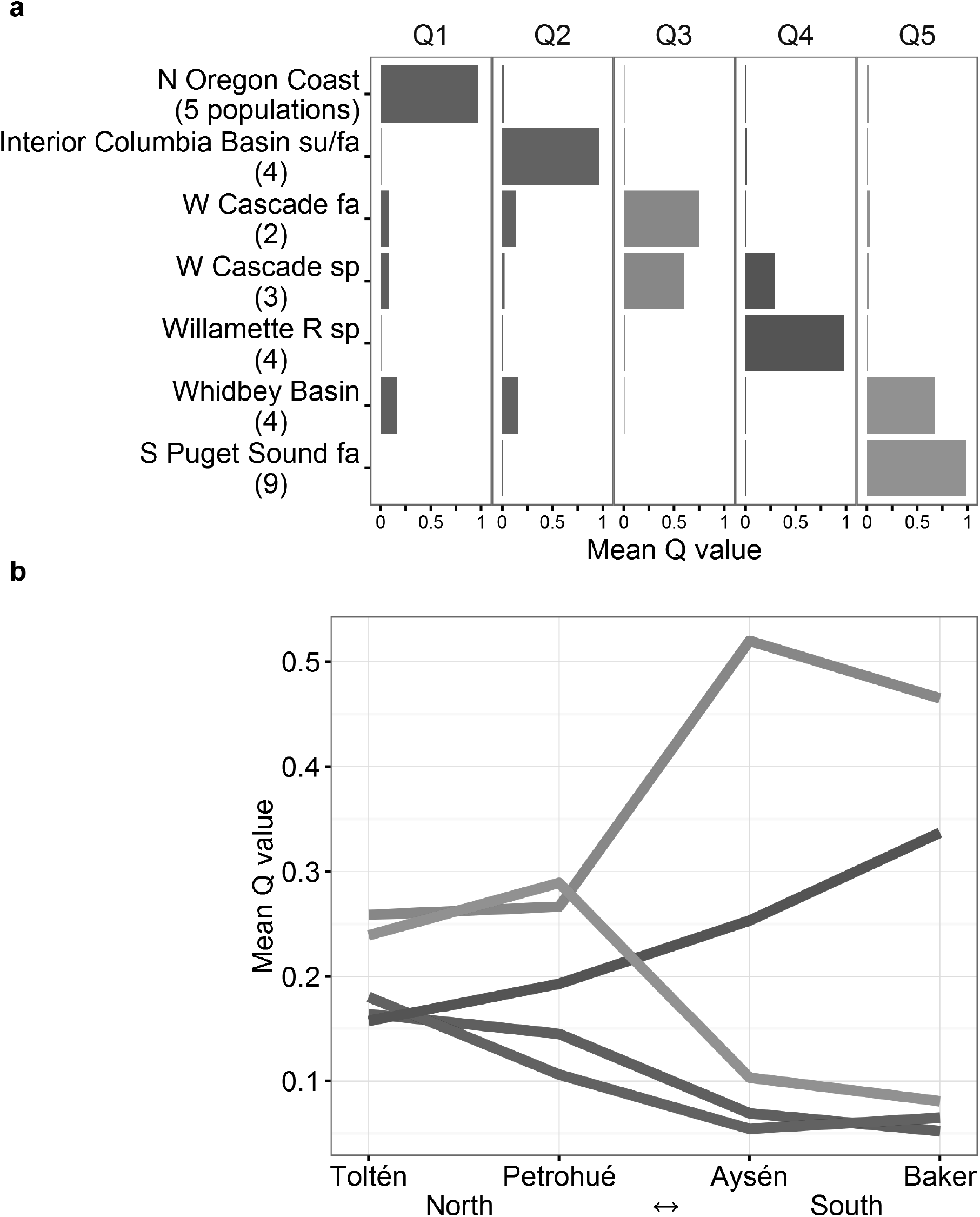
Latitudinal cline in lineage diversity as inferred from model-based clustering. The clearest discrimination among the 31 baseline populations selected for this analysis was achieved at five clusters (K = 5), which associated (Q1-Q5) to known North American lineages (a). Membership of Patagonian samples to these clusters (i.e., inferred ancestry), showed a decline in lineage diversity from north to south, consistent with CML mixture analysis. Standard ΕK plot, and STRUCTURE plots, are available in Supplementary Fig. S3 and Fig. S4.

## Discussion

We investigated the phylogenetic ancestry of introduced Chinook salmon in Patagonia by conducting two different classes of genetic mixture analysis. Diverse genotypes led to the identification of many putative ancestral sources introduced primarily from the states of Oregon and Washington. Our results were largely consistent with historical records of fish introductions and recent molecular genetic studies. However, we also found interesting differences such as apparent contributions from undocumented introductions, and conversely, lack of evidence supporting the naturalization of well-documented introductions.

Our study is distinguished from previous molecular studies of Patagonian Chinook salmon (^16,19,29,30^) in three principal ways: 1) We used the most inclusive baseline dataset possible, including all potential North American donor lineages. Previous studies have been more or less limited by the number of reference lineages available for particular genetic markers. The GAPS microsatellite baseline is the most extensive of its kind and is ideally suited for this application. 2) To our knowledge, ours is the first study of its kind to take into account the potential for spurious allocation of admixed genotypes. Other studies have addressed genetic drift in introduced populations ^29^, but not admixture between divergent lineages. We suggest that admixture may be more problematic than genetic drift, and simulations of mixed-origin founding offered a cautionary note on potentially spurious results. Our observations are relevant to many studies that seek to characterize small genetic contributions from multiple founding lineages ^36^. 3) Our study included naturalized populations both within and outside salmon aquaculture zones. This helped clarify the poorly documented role of net-pen aquaculture on Chinook salmon introduction to Chilean Patagonia. Thus, our study provides important new insight into the ancestry and history of colonization of Patagonian Chinook salmon.

Analyses of allele frequencies led us to pool within-river-basin samples into putative populations, yet additional evidence suggested substructure, or non-equilibrium conditions, within river basins. Because our samples were small it was important to evaluate the strength of assuming basin-level population structure from different perspectives, in addition to analyses of allele frequencies. Thus, we also tested for departures from expected Hardy-Weinberg genotypic proportions, especially heterozygote deficits that might indicate a Wahlund Effect resulting from sampling two or more genetically distinct populations. This analysis would help us evaluate population structure and potential assortative mating among sympatric founding lineages. Mating structure is critical to our understanding of effective population size and potential for adaptation and persistence of introduced lineages in Patagonia. We did find elevated *F*^IS^ values within basins (putative populations), consistent with assortative mating or other non-equilibrium conditions, but the *F*_IS_ estimates were not significantly greater than zero. Recognizing limited power for detecting a Wahlund Effect, we also tried to draw inference from CML individual assignment probabilities. Fewer intermediate assignments (split probabilities between reference groups) were observed than expected for a (simulated) mixed-origin population at equilibrium. If real, such a result might be consistent with, for example, assortative mating of two or more genetically distinct lineages within a river basin. Alternatively, strong out-breeding depression ^e.g., ,37^ or insufficient time since introductions could result in fewer than expected mixed-origin individuals. Resolution of this question will require larger sample sizes within and among sites and river basins as well as across multiple generations. Alternatively or additionally, more markers could be surveyed. For example, genome sequencing would provide haplotype arrays that might be quite powerful for evaluating introgression and equilibrium.

Diversity of founding lineages was higher in northern Patagonia, and yet this latitudinal trend was uncorrelated with standard measures of genetic diversity. Overall, population genetic diversity measured by heterozygosity and allelic richness was higher than expected in Patagonia— nearly identical to North American populations. Our genetic CML mixture analyses also suggested a diverse array of founding lineages in Patagonia. Initially, we assumed high genetic diversity, i.e., heterozygosity and allelic richness, was a result of mixed ancestry. Previous studies have made similar conclusions ^19,29,30^. CML mixture analysis and M-BC both estimated increasing diversity of founding lineages from south to north in Patagonia. Surprisingly, however, populations with the most diverse ancestry showed no higher levels of heterozygosity or allelic richness. Nor was there a spatial gradient for heterozygosity or allelic diversity, as was evident for lineage diversity.

It would seem almost axiomatic that a diversity of founders would introduce more alleles relative to monophyletic populations, yet other factors might be confounding a clear association between lineage diversity and genetic diversity. Given our broad and overlapping estimates of heterozygosity and allelic richness, it might simply be that our sample sizes were too small to obtain a precise view of genetic diversity relative to lineage diversity.

A principal challenge in this study was to distinguish the genetic signal of true ancestry from the noise created by mixed-origin, introgressed populations. In our CML mixture analysis we expected some misassignments between related populations. The patterns of misassignment in the North American reference baseline were well understood based on previous results ^27^. However, our simulation of mixed-origin founding revealed some spurious genetic allocation not previously reported in studies that used this approach ^29,30,36^. In the empirical results, we observed allocations of up to 6% estimated genetic contribution from very unlikely donor regions, such as Nass River and the South Thompson River, both in British Columbia. This 6% level of contribution is equivalent to a little more than one fish with a high assignment probability in a population sample of the size we collected. The CML simulation showed us that mixed origin (admixed) individuals usually assign to the true donors, however, some individuals will assign to unrelated populations, sometimes with high assignment probability. Note, however, that the probability values are conditional on available baseline references. A high assignment probability might simply mean that no other reference population could likely have produced a given genotype. This is relevant, for example, to studies that assume a probability threshold (e.g., 0.8) for accurate assignment, and especially to those with incomplete baseline datasets. While a probability threshold such as 0.8 is clearly useful in conventional genetic mixture applications ^28^, in a mixed-origin founding application, it is almost certainly the case that some high-probability individual assignments are attributable to the misassignment phenomenon that we demonstrated through simulation. Apparently, introgression between donor populations results in unique genotypes that assign arbitrarily to other, unrelated populations. To avoid falsely implicating potential donor lineages we set a lower threshold for genetic contribution below which putative ancestry was viewed with caution and some skepticism. Based on review of both simulated and empirical data, we used a threshold of 10% estimated genetic contribution to any single population in Patagonia. It was not that we rejected as potential contributors lineages below 10%, rather we were less confident in identifying those as founders.

Despite this caveat, our findings clearly confirm the importance of entrepreneurial attempts to establish Chinook salmon open-ocean ranching programs in Chile during the late 1970s and 1980s. In particular, the West Cascade spring and fall lineages, introduced from the Lower Columbia River, were likely associated with the enterprises suspected to have initiated the invasion. In 1978 Domsea Pesquera Chile Ltda. (a subsidiary of Union Carbide Corporation, USA) began yearly stockings of Chinook salmon in a small stream on Quinchao Island, Curaco de Vélez, Chiloé (42°S). Three strains of Chinook salmon were introduced: Cowlitz River spring run, Bonneville Hatchery fall run, and University of Washington Hatchery fall run, in decreasing number of individuals released ^21,38^. The primary stock that appeared to become established was Cowlitz River spring run ^20,38,39^; precisely the same primary ancestral lineage we identified in our samples. The finding of the related fall-run lineage (West Cascade fall) in our samples might correspond to the Bonneville Hatchery fall-run, whereas there is little evidence of successful University of Washington Hatchery fall-run (see below).

Regarding the introductions in Río Chirri (Río Bueno) in the early 1970s from Green River Hatchery (also located on the Cowlitz River), establishment seems unlikely. Although this population is from the West Cascade fall lineage, to our knowledge there are no records of Chinook salmon in Río Bueno during the decade that followed introduction ^18^, and no fish in our analyses assigned to Green River Hatchery. Nevertheless, definite confirmation would require analyses of samples from Río Bueno.

Subsequent open-ocean ranching experiments in the Magellan region (51°S) utilized the University of Washington Hatchery fall-run stock ^21,22^. Here, Salmones Antártica S.A., initiated stockings first in Río Santa María (1982) and then in Río Prat (1983-1989) using University of Washington Hatchery fall-run and the newly established Chilean-based brood stocks (Table 1). Nevertheless, we found little evidence supporting the successful colonization by University of Washington Hatchery fall-run stock introduced in Patagonia. This stock, also known as Portage Bay fall-run, from the South Puget Sound fall reporting group, was primarily derived from the Green River draining to Puget Sound (different to Green River in the West Cascades), particularly from Soos Creek Hatchery (1949-1950s), though exchanges with other populations took place over the years ^40^. The Soos Creek Hatchery itself had exchanges with many other populations, mainly within the Puget Sound area. It is puzzling that in our study not a single fish showed any probability of assignment to the University of Washington reference population. Only five fish (17% genetic contribution) from Petrohué River assigned to Soos Creek Hatchery.

Previous studies have inferred ancestral affiliation from the University of Washington Hatchery based on mitochondrial DNA ^16,19^, but owing to the low resolution of those analyses (incomplete baselines; geographically wide-spread haplotypes), it is unclear if those were real signals. Other investigators, using SNPs and a more extensive baseline ^41^, found little evidence to support South Puget Sound ancestry, although they still suggested a possible contribution of University of Washington Hatchery stock based on its supposed origin from lower Columbia River lineages, which do appear as major contributors to Patagonian populations ^29,30^. Our results provided no support whatever for that interpretation. The GAPS baseline shows no similarity between Lower Columbia River populations and University of Washington Hatchery stock—or indeed any other Puget Sound populations. University of Washington Hatchery brood stock is genetically distinct with 80% correct assignment of known-origin fish in leave-one-out jackknife analysis. All University of Washington Hatchery fish assign correctly to the South Puget Sound fall run lineage (i.e., no misassignments to West Cascade spring nor to any other population outside South Puget Sound).

Given the historical record and our genetic mixture results, we suggest three possible outcomes for the introductions from University of Washington Hatchery fall-run stock into Patagonia: (i) Naturalization failed, leaving no genetic traces in current Patagonian populations. (ii) Naturalization succeeded to some extent, and assignments to South Puget Sound fall represent remnant evidence (through admixture). (iii) Naturalization succeeded in the Magellan region, yet subsequent range expansion did not reach our study area, or those of other studies in Argentinean and Chilean Patagonia ^29,30^. In any case, there is little doubt that open-ocean ranching experimental operations in Chile in the late 1970s and 1980s were responsible for the introduction of West Cascade spring Chinook salmon in Patagonia.

Patagonian Chinook salmon unrelated to the West Cascade spring and fall lineages (hereafter non-WC) accounted for almost half of the fish (45.8%), mostly taken from north of parallel 45°S. The phylogenetic origins of non-WC fish were tracked to 14 diverse and geographically widespread lineages in North America. This seemingly hyperdiverse assemblage is at least partly explained by spurious individual assignments related to mixed ancestry (consistent with our simulations). However, much of this diversity appears due to accidental escapes from thriving Chilean net-pen salmonid aquaculture operations in the 1990s that imported ova from diverse sources. Since 1988, Chilean salmonid aquaculture shifted entirely to net-pen aquaculture. Although substantial escapes were frequent, they were hard to quantify because they commonly went unreported ^42–45^. Data gathered from insurance companies alone ^46^ led to an estimate average of one million salmon and trout escaping every year, primarily from net-pen aquaculture. Although Chinook salmon farming has always been marginal in Chile relative to other species (<4,000 tons y^−1^ and <2.5% y^−1^ of total salmonid production ^47^), sustained Chinook salmon farming in leaky rearing facilities certainly contributed propagules to the wild ^42^. We estimated that with one million escaped salmon and trout per year, adjusting for the contribution of Chinook salmon to annual harvest ^47^, *c.* 100,000 farmed sub-adult Chinook salmon might have escaped between the onset of modern salmonid farming (1988) and the date of our sampling (early 2004). This conservative estimate illustrates the potential of net-pen aquaculture as a major source of colonists. During this period, numerous national and trans-national aquaculture companies were established in Chile, propagating Chinook salmon brood stocks to meet increasing demand. Aedo ^24^ assembled the fragmentary record of importations and concluded that ova originated at least from Washington, Oregon, Vancouver Island, and New Zealand (of Central Valley ancestry), but specific stocks mostly went undocumented (Table 1).

Although sparse, this information could explain the incidence of Patagonian genotypes related to multiple lineages of the West Coast of U.S.A. Furthermore, the spatial distribution of non-WC fish was concentrated in the north (Toltén and Petrohué) and is consistent with more intensive propagule pressure of distinct lineages farmed in this region. In fact, during the 1990s, most aquaculture activity (and escapes) was concentrated in the Lakes District region (41-43°S), within 200 km of the Petrohué River. Toltén River (39°S), being a few hundred kilometres further north, might have received colonists dispersing naturally from the Lake District region, escapees from inland hatcheries, or deliberate stockings, because this is a well-populated region easily accessible by paved roads. In sharp contrast, in the more remote south of our study area (45-50°S), aquaculture activity was only incipient (Aysén River) or non-existent (Baker River). Here, accidental escapes or unreported deliberate stockings were less likely, and natural dispersion alone seems the most likely source of founders, especially in the remote and isolated Baker River basin. Other studies have come to similar conclusions by comparing one Chinook salmon population in the Pacific slope of Patagonia (Futaleufú River, 43°S ^30^) to another population in the Atlantic slope (Santa Cruz River, 50°S ^29^). The population closer to the Lake District region (Pacific slope) showed affinity to more lineages than the population in the Atlantic slope ^30^. Thus, escapes from net-pen aquaculture activity in the Lake District region have augmented the genetic diversity initially introduced by open-ocean ranching operations ^19,30,this study^. Similarly, the introgression of cultured rainbow trout genes into pre-existing, naturalized rainbow trout populations in Patagonia, has also been demonstrated ^48^.

An apparent lack of Central Valley ancestry was similar to the findings of one previous molecular study ^29^ but different to another ^30^. It is worth noting that previous studies like ours did not account for potential spurious assignments of mixed-origin fish to unrelated reporting groups. Thus, numerous additional sources were identified as putative contributing lineages. We do not refute the presence of these additional lineages. We simply recognize the limitation we discovered in our genetic mixture analysis regarding mixed-origin fish, and we focus on what we take to be the major lineages that became established at our study sites (those lineages receiving ≥ 10% allocation).

Notable patterns of lineage distribution are emerging at a continental scale. Northern study sites (Toltén and Petrohué) showed unexpectedly high lineage diversity, including substantial estimated contributions from five or more lineages (North Oregon Coast, West Cascade spring, South Puget Sound fall, Whidbey Basin, and Interior Columbia River summer/fall). By contrast, southern study sites (Aysén and Baker) revealed a striking dominance of three closely related lineages from the Lower Columbia River (i.e., West Cascade spring, West Cascade fall and Willamette spring lineages). A similar diversity was also found in the Santa Cruz River draining to the Atlantic Ocean in Argentinian Patagonia ^29^. A number of possibilities could explain this interesting pattern: (1) As discussed above, higher propagule pressure of distinct lineages leaking from fish farms might have resulted in higher lineage diversity in northern sites compared to remote southern localities (propagule pressure hypothesis). (2) Most dispersing colonists moved northwards in our study area such that those colonizing Aysén and Baker rivers were derived from stockings in the Magellan region of West Cascade spring/fall ancestry, whereas those colonizing Petrohué and Toltén rivers were derived from fish farms in the Lake District region (propagule+dispersion hypothesis). This pattern would resemble natural ocean migration patterns of Washington and Oregon stocks towards the north in the North Pacific, although it is unclear how the fish would respond to magnetic stimuli of the southern hemisphere ^49^. (3) Deliberate stockings initiated the first wave of invasion that eventually reached Aysén and Baker rivers, either from the north or south. Previous establishment of West Cascade spring/fall lineages might have inhibited the expansion of farmed lineages from the north (density-dependent hypothesis) ^50–52^. At present, it is hard to gauge the relative importance of these processes, and it is conceivable that each contributed to the observed pattern of lineage distribution. Future work should increase the geographic coverage, number of locations and sample sizes, and should incorporate time-series to study the consequences of anthropogenic sequential introductions, dynamics of range expansion, and lineage mixing and admixture.

Complex invasion dynamics and distinct patterns of genetic diversity are likely to have profound evolutionary consequences in the continuing range expansion of Chinook salmon in Patagonia. High genetic diversity, as measured by high allelic and lineage richness ^30, this study^, high phenotypic plasticity and rapid adaptive evolution observed in Chinook salmon introduced to New Zealand ^11,15^, as well as pre-adaptation to environmental conditions inferred for Patagonian populations ^18^, are all features that suggest high adaptive potential of Chinook salmon in Patagonia. Lower-than-expected levels of heterozygosity, if confirmed and maintained over generations, could indicate reproductive isolation between sympatric lineages, which would help maintain lineage identity.

Whether this happens is crucial to the evolutionary trajectory of the species in its new range. Lineages could either evolve adapting genetically to local conditions ^15^ or they could interbreed and recombine, potentially resulting in new phenotypic variation with unpredictable evolutionary outcomes. Chinook salmon may exert strong ecological impacts in freshwater, estuarine and marine ecosystems ^18^, and different evolutionary outcomes will directly affect these impacts owing to considerable phenotypic variation among lineages.

## Materials and methods

### Field collection

Chinook salmon populations were sampled in the fall of 2004 from rivers in the four Andean watersheds of Toltén, Petrohué, Aysén and Baker (39-48°S), all draining to the South Pacific through Chile (Figure 1). The results presented here are from 87 fish (after exclusion of 9 fish for which DNA could not be extracted or genotyped). Adult spawners made up 91% of our sample (weight range 2250-21000g), and 9% were juvenile parr (9-37g). Collection took place near spawning habitat using a variety of methods including gillnets, hooks, hand-line, and fly-fishing. A fin clip was dissected from each fish and preserved in 96% ethanol for later genetic analyses. A more detailed description of the sampling and sites is available elsewhere ^18^.

### Population genetic analysis

DNA was extracted from individual fish tissue samples by using silica-based purification (QIAGEN DNeasy 96 Blood and Tissue Kit, rat tail protocol). Genomic DNA samples were amplified for 13 highly polymorphic microsatellite loci and genotyped by using an Applied Biosystems Incorporated (ABI) 3100 Genetic Analyzer and ABI’s Genescan and Genotyper software (v3.7), according to methods of Van Doornik et al. ^53^. We used the GAPS Chinook salmon microsatellite markers following the GAPS consortium conventions for standardization of genotypic data ^25^. We calculated allelic and genotypic frequencies as well as diversity metrics allelic richness, *AR* (based on minimum sample size of 12 individuals), and expected heterozygosity, *H*_S_, estimated with FSTAT v2.9.3 (Goudet 2001). Fixation indices *F*_ST_ and *F*_IS_ were represented by Weir and Cockerham’s ^54^ multilocus estimators, *θ* and *f*, calculated with FSTAT v2.9.3 ^55^. *F*_ST_ was used to evaluate relationships among populations (see below) whereas *F*_IS_ was used to evaluate departures from Hardy-Weinberg-Castle (HWC) expectations for genotypic proportions within putative populations ^56^. We also tested for composite linkage disequilibrium by using the Markov chain Monte Carlo (MCMC) method implemented in the program GENEPOP v4.2 ^57,58^. Before proceeding with our population genetic analyses, we tested for loci that departed from neutral expectation in order to avoid loci that might bias our parameter estimates and genetic distance estimates (although departures from neutrality would not necessarily bias our genetic mixture analysis). We used the *F*_ST_ outlier approach ^59^ implemented in the LOSITAN software package ^60^. Genetic differences among the South American river basins were analyzed in more detail utilizing pair wise *F*_ST_ values and leave-one-out jackknife self assignment among introduced populations in order to characterize genetic differences among the introduced Patagonian populations. Because our sample sizes were small, we were especially cautious of non-significant results because we had little power to reject null hypotheses, even when false. However, where we did observe statistical significance, we generally trusted our results as biologically meaningful, despite small sample sizes.

### North American baseline dataset

We relied on a baseline dataset of North American reference populations that were selected from the GAPS baseline ^25^ to represent as closely as possible historical phylogeographic lineages in North American Chinook salmon. Specifically, we used a slight modification of the dataset analyzed by Moran et al. ^32^. We added the following populations because they were identified previously as potential sources of Patagonian Chinook salmon (Table 1): Cowlitz River spring and fall-run, and Green River fall-run from the West Cascade spring and fall-run reporting groups, and University of Washington Hatchery fall-run population from the South Puget Sound fall-run reporting group.

Reporting groups are intended to reflect phylogenetic lineages, and here we use either terminology depending on whether the emphasis is on methodology (reporting groups) or phylogeography (lineages). In total, our baseline dataset consisted of 19,973 individuals from 146 populations nested within 46 reporting groups distributed from Central Valley, California (40°N) to the Yukon River in British Columbia (64°N) (Figure 1). The baseline is widely accepted in the Pacific salmon genetics community as being both comprehensive and temporally stable ^25^. Even very small headwater populations show long-term stability of allele frequencies ^53^.

### Conditional maximum likelihood mixture analysis

Two different methods were used to explore the likely ancestry of introduced Chinook salmon populations in Chile, conditional maximum likelihood (CML) and model-based clustering (M-BC). As a first approximation of genetic similarity and potential ancestry, we used the Rannala and Mountain ^31^ CML algorithm implemented in the ONCOR software package ^61^. That analysis allowed us to estimate the posterior probability distribution for baseline population membership of each individual fish collected in Patagonia. Individual fish were assigned to the population for which they had the maximum *a posteriori* probability of membership. Results were subsequently aggregated by reporting group (genetic lineage), which is the level of inference in essentially all our interpretations. The mean probability across individuals for membership to a particular lineage was taken as an approximation of overall genetic ancestral contribution to the population (normally interpreted as the unbiased estimate of that population’s contribution to, e.g., a mixed-stock fishery). Simulation studies helped us evaluate the strength of those assumptions under conditions of mixed ancestry. The CML analysis was used as a first approximation to help narrow the range of contributing lineages to just the major contributors. A reduced set of populations and lineages was then further analyzed by M-BC.

### Simulated mixed-origin founding

Despite the common use of CML for studies of ancestry of non-native populations ^e.g., ,29,30,36^, its validity in that application has not to our knowledge been demonstrated. As a test of the algorithm’s sensitivity to the presence of mixed-origin founding, we simulated a new population derived from large and equal numbers of individuals from two of our North American baseline populations. These were specific populations that appeared to be associated with the introduction to Patagonia (Cowlitz River Hatchery spring-run population in the West Cascade spring-run lineage and Soos Creek Hatchery fall-run population in the South Puget Sound fall-run lineage). That simulated founding involved the creation of an ideal population (at mutation/drift equilibrium) with intermediate allele frequencies to the Cowlitz River and Puget Sound source populations. We then drew 1000 individuals from that new, mixed-origin population, and we estimated the posterior probability distributions for those simulated individuals, the same as we did for the real Patagonian fish. We sought to determine whether the admixed genotypes would assign with high probability to one or the other source population, or split their probability of assignment between the two source populations. These novel genotypes might even assign to other, unrelated populations in the large coast-wide North American baseline (where “assignment” simply reflects the maximum *a posteriori* probability of membership to population or reporting group). We needed to know the extent to which potentially spurious assignments might affect our analysis of the Patagonian populations. As a benchmark for misassignment in the North American reference populations, we conducted a leave-one-out, jackknife analysis for comparison of *a posteriori* probability distributions between simulated mixed-origin fish and real fish, from both North and South America (Supplementary Figure 2).

### Model-based clustering

Based on the results of our CML mixture analysis we conducted a more focused model-based cluster analysis (M-BC) of the introduced fish relative to the putative source populations using computer program Structure v2.3.4 ^33^. From the North American reference baseline described above, we selected 8,228 fish from 31 populations distributed among seven lineages to include in our analysis. Selected North American lineages for M-BC included the following: Whidbey Basin, S. Puget Sound fall, Interior Columbia Basin summer/fall, Willamette River spring, West Cascade fall, West Cascade spring, and North Oregon Coast. The North American populations and lineages selected for M-BC were those identified in CML as major contributors by virtue of *c.* 10% allocation or higher to any single Patagonian population (North Oregon Coast was included because allocation to that lineage was quite close to our threshold, rounding up to 10%. Including North Oregon Coast as a possible source seemed to be conservative, especially with records of potential introductions from Northern Oregon). The 10% value was selected based on an apparent break in the composition estimates. We also inferred from simulations that allocations of less than 10% might be due at least in part to spurious assignment of mixed-origin fish (see *Methods: Simulated mixed-origin founding*).

For the M-BC analysis, we conducted 110,000 MCMC realizations per chain, discarding the first 10,000 iterations as a burn in. Apparent convergence of diagnostic parameters was observed within the first 10,000 iterations (i.e., α, *F*, divergence among populations *D*_i,j_, and the likelihood estimate). We used an admixture model, including location information ^62^, with allele frequencies correlated among populations ^63^. We assumed population specific *F*_ST_ values, and updated allele frequencies by using baseline individuals only, thus treating Patagonian samples as having unknown origins. Multiple MCMC chains (10) were constructed for each value of *K* (number of ancestral clusters). We modeled values of *K* from two to nine for the North American populations. The most appropriate *K* value was selected based on likelihood and ∆*K* ^34,64^, as well as optimal discrimination of putative North American source populations. Structure output was processed with the STRUCTURE HARVESTER computer program ^35^, and plotted in the R computing environment version 3.3.1 ^65^.

## Acknowledgements

We especially thank Mart Gross for his pivotal support at the initial stages of this study. Comments by Mart Gross, Jim Myers, David Teel, Robin Waples, and Jeff Hard helped us improve earlier drafts of this article. Jon Wittouck kindly provided historical records of fish transfers from the University of Washington Hatchery. Cristián Correa was funded by the Government of Chile trough grants CONICYT-PAI N°82130009, and FONDECYT-Iniciación en la Investigación N°11150990.

## Author contributions

C.C conceived the study, and collected samples from Patagonia. P.M conducted the genetic analyses. C.C prepared all figures and tables, and both authors contributed to the interpretation of results and writing of the manuscript.

## Additional information

The authors declare no competing financial interests.

## Supplementary Information

**Table S1:**
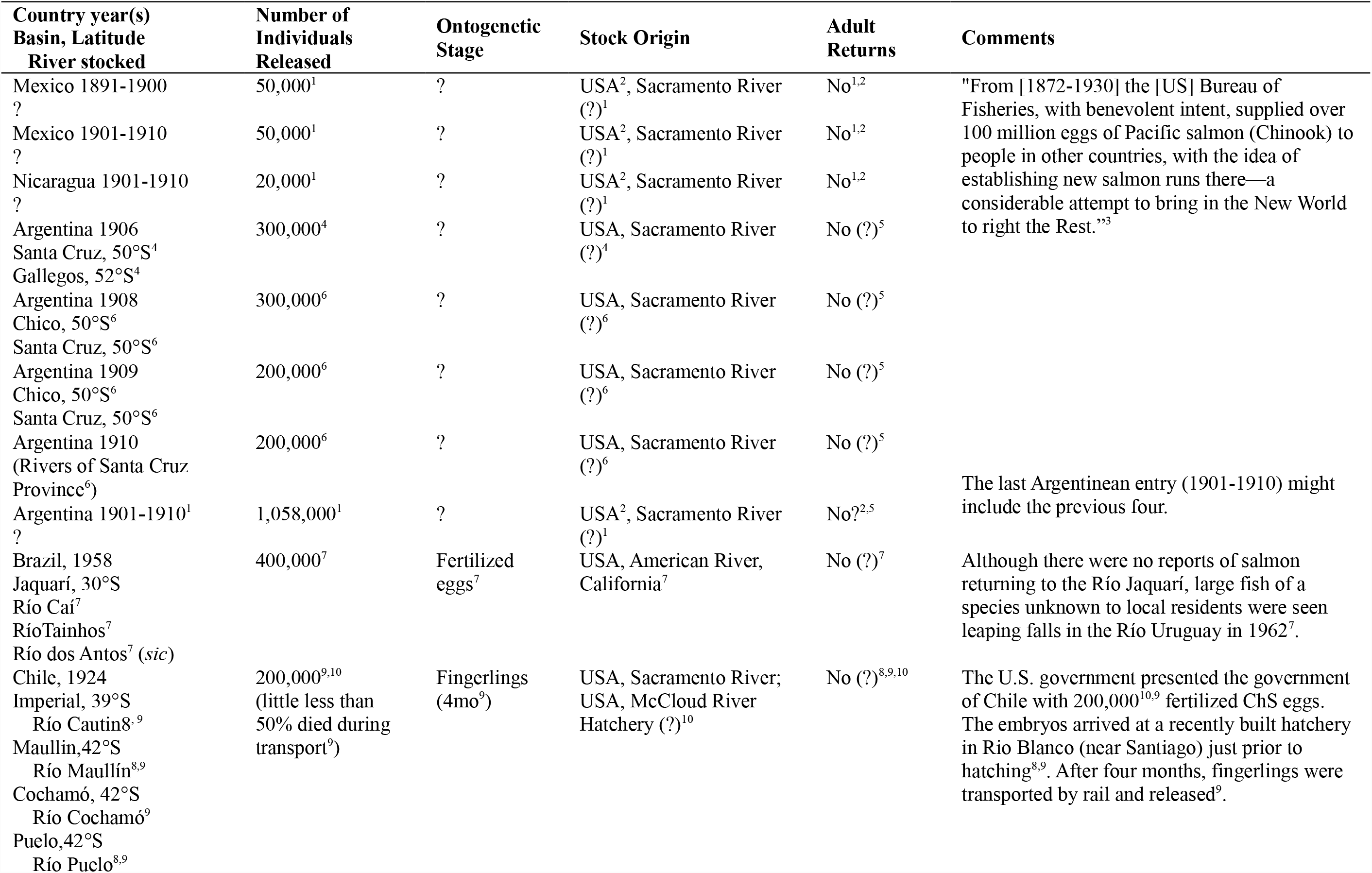
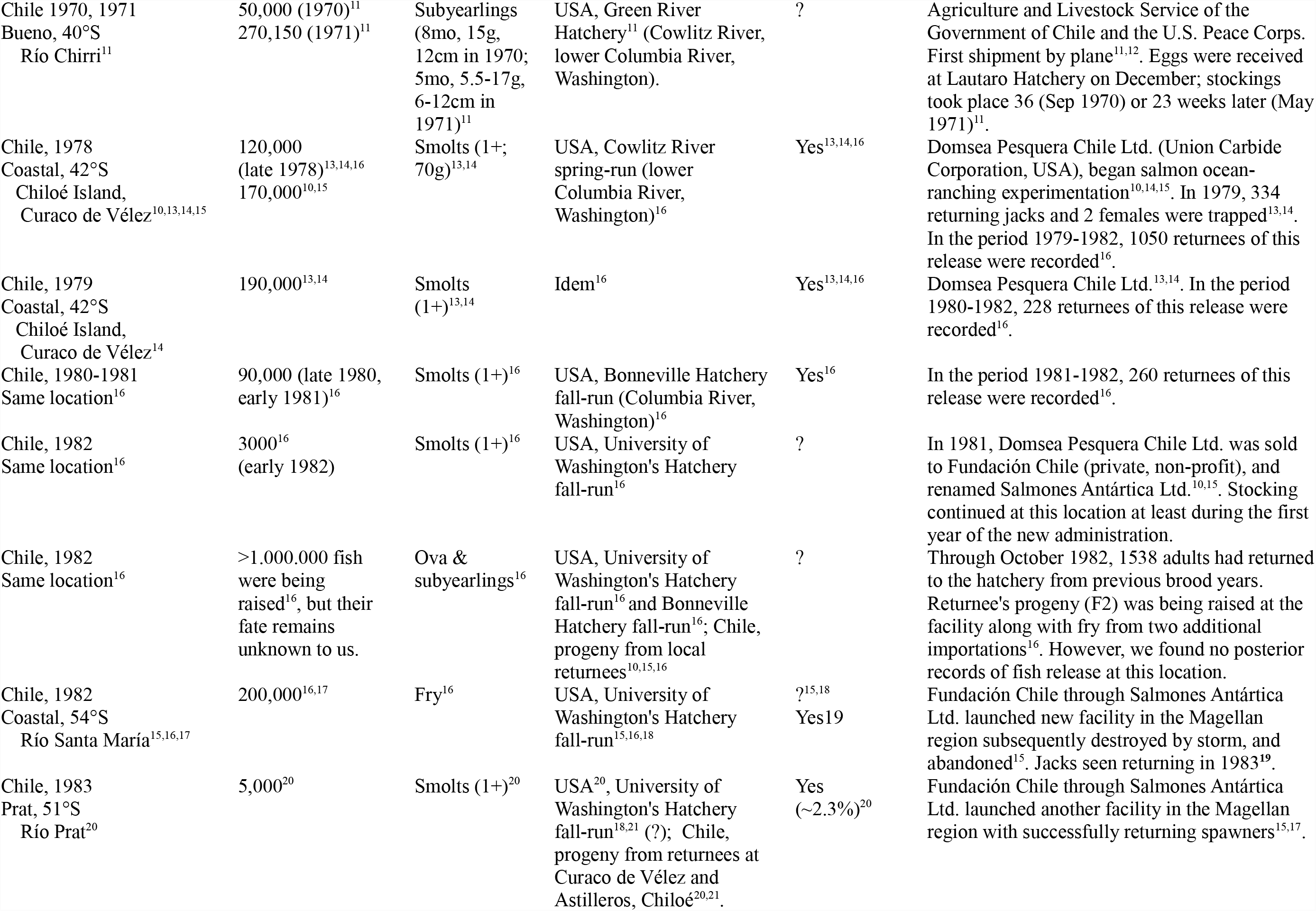
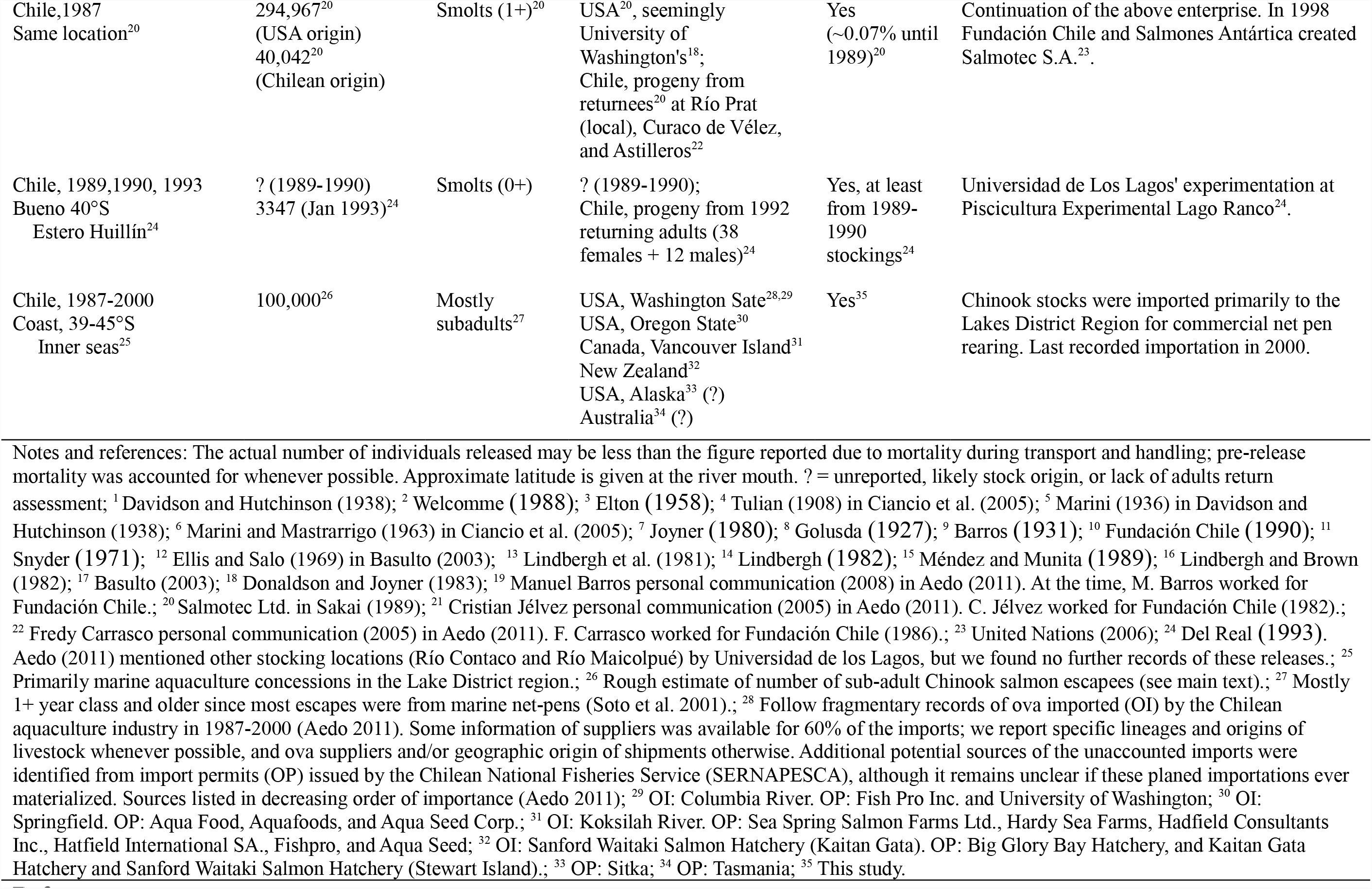
Chinook salmon deliberate and accidental releases in Latin America (modified from Correa and Gross 2008).

**Table S2:**
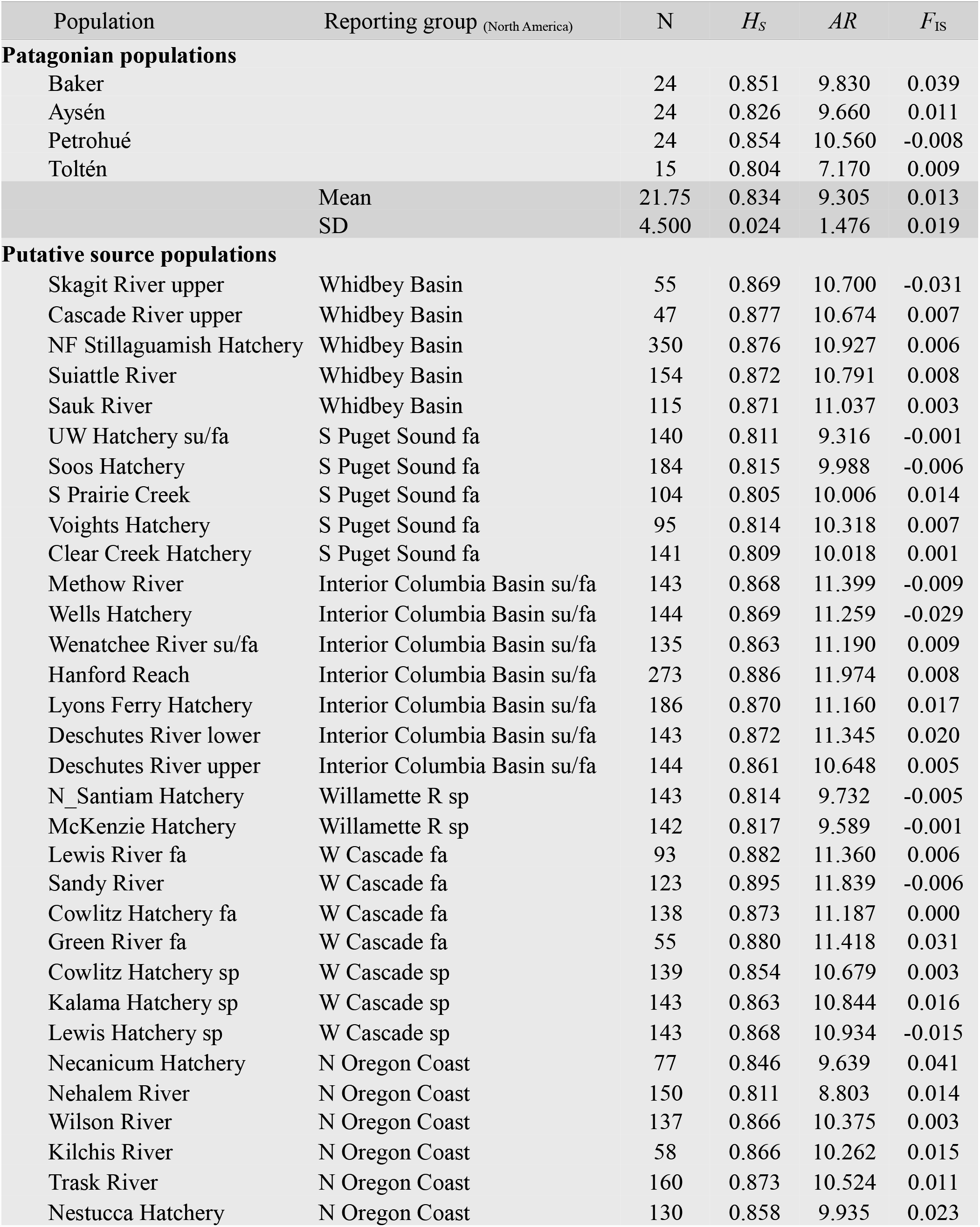
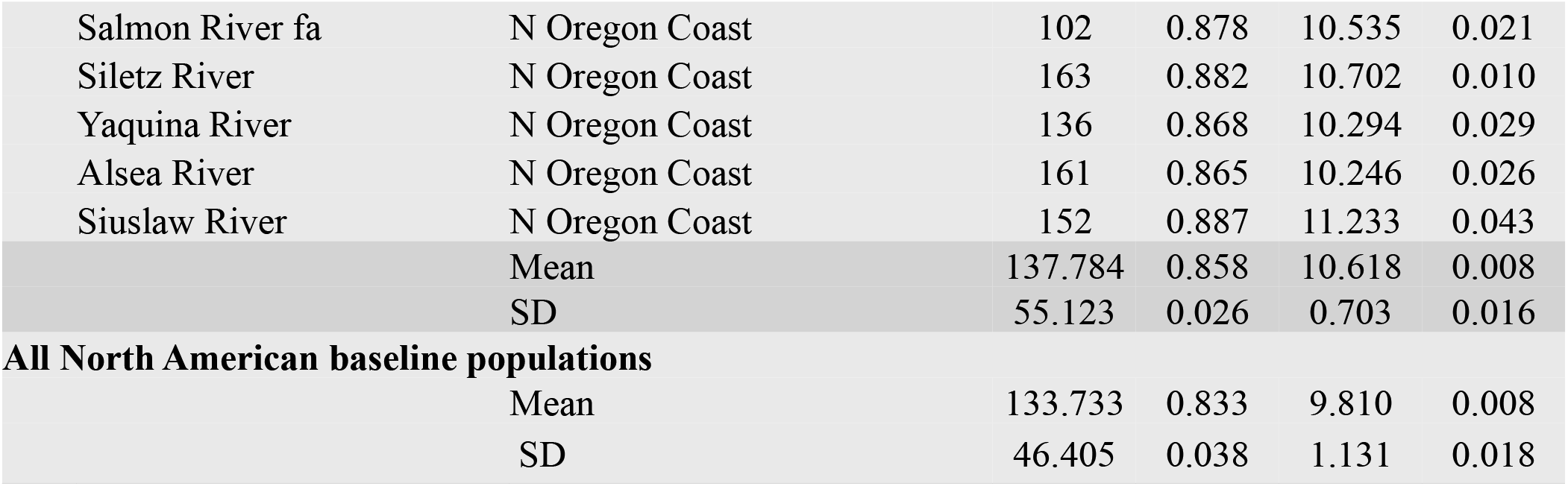
Genetic diversity in Patagonian populations, putative North American founding populations, and overall North American baseline populations.

**Table S3:** Genetic ancestral contribution of North American lineages to Patagonian Chinook salmon based on population-level CML mixture analysis.

| ID | Reporting Group | NA Population | Patagonian watershed |  |  |  | Pooled |
| --- | --- | --- | --- | --- | --- | --- | --- |
|  |  |  | Toltén | Petrohué | Aysén | Baker |  |
| 1 | Central Valley fa | Stanislaus R |  | 4.1 (1) |  |  | 1.2 (1) |
| 1 | Central Valley fa | Tuolumne R |  | 4.1 (1) |  |  | 1.2 (1) |
| 5 | Klamath R | Klamath R fa | 6.9 (1) |  |  |  | 1.3 (1) |
| 6 | Chetco R | Chetco R | 7.3 (1) |  |  |  | 1.3 (1) |
| 9 | Willamette R sp | N Santiam H |  |  | 4 (1) | 3.3 (1) | 1.9 (2) |
| 9 | Willamette R sp | McKenzie H |  |  | 1.3 (0) | 15.0 (4) | 4.6 (4) |
| 13 | N Oregon Coast | Salmon R f | 0.6 (0) | 4.1 (1) |  |  | 1.3 (1) |
| 13 | N Oregon Coast | Siuslaw R | 2.9 (0) | 4.2 (1) |  | 0.5 (0) | 1.9 (1) |
| 13 | N Oregon Coast | Trask R | 5.9 (1) | 0.1 (0) |  |  | 1.1 (1) |
| 15 | W Cascade fa | Cowlitz H fa | 8.9 (2) | 3.2 (1) | 16.8 (3) | 5.4 (1) | 8.2 (7) |
| 15 | W Cascade fa | Sandy R | 0.6 (0) | 0.6 (0) | 0.5 (0) | 5.5 (2) | 2.0 (2) |
| 16 | W Cascade sp | Kalama H sp | 3.0 (0) | 15.2 (4) | 26.9 (7) | 28.7 (7) | 19.7 (18) |
| 16 | W Cascade sp | Cowlitz H sp | 4.3 (1) | 6.4 (1) | 44.4 (8) | 35.8 (8) | 23.7 (18) |
| 17 | Interior Columbia Basin su/fa | Wenatchee R s/f | 6.8 (1) | 0.4 (0) |  |  | 1.4 (1) |
| 17 | Interior Columbia Basin su/fa | Hanford Reach | 6.2 (1) | 8.5 (2) | 4.4 (1) | 5.5 (1) | 6.2 (5) |
| 17 | Interior Columbia Basin su/fa | Lyons Ferry H | 6.0 (1) | 0.1 (0) |  | 0.1 (0) | 1.1 (1) |
| 19 | S Puget Sound fa | Clear Cr H | 0.1 (0) | 2.1 (0) | 1.3 (0) |  | 0.9 (0) |
| 19 | S Puget Sound fa | Soos H |  | 16.9 (5) |  | 0.1 (0) | 4.9 (5) |
| 19 | S Puget Sound fa | S Prairie Cr | 0.1 (0) | 1.0 (0) |  |  | 0.3 (0) |
| 22 | Washington Coast | Sol Duc H |  | 4.0 (1) | 0.1 (0) |  | 1.2 (1) |
| 22 | Washington Coast | Forks Cr H | 6.5 (1) | 4.0 (1) | 0.1 (0) |  | 2.4 (2) |
| 23 | Straits of Juan de Fuca | Elwha R | 0.2 (0) | 5.7 (1) |  |  | 1.7 (1) |
| 24 | Whidbey Basin | Suiattle R |  | 4.2 (1) |  |  | 1.2 (1) |
| 24 | Whidbey Basin | Cascade R U | 20.1 (3) | 0.3 (0) | 0.2 (0) |  | 3.8 (3) |
| 26 | E Vancouver Is | Big Qual H | 6.3 (1) | 2.2 (1) |  |  | 1.8 (2) |
| 31 | S Thompson R | L Adams H | 6.4 (1) | 0.1 (0) |  |  | 1.2 (1) |
| 38 | SSE Alaska | Clear Cr | 0.6 (0) | 4.4 (1) |  |  | 1.4 (1) |
| 39 | Nass R | Kincolith R |  | 3.9 (1) |  |  | 1.1 (1) |
| No. Individuals in the mixture |  |  | (15) | (24) | (20) | (24) | (83) |

Notes: Values represent average percent genetic contribution; in brackets, frequency of individual assignments to baseline populations, as inferred from individual's highest assignment probability. Identifiers (ID) correspond to those in Figure 1 (main article).

**Figure S1:**
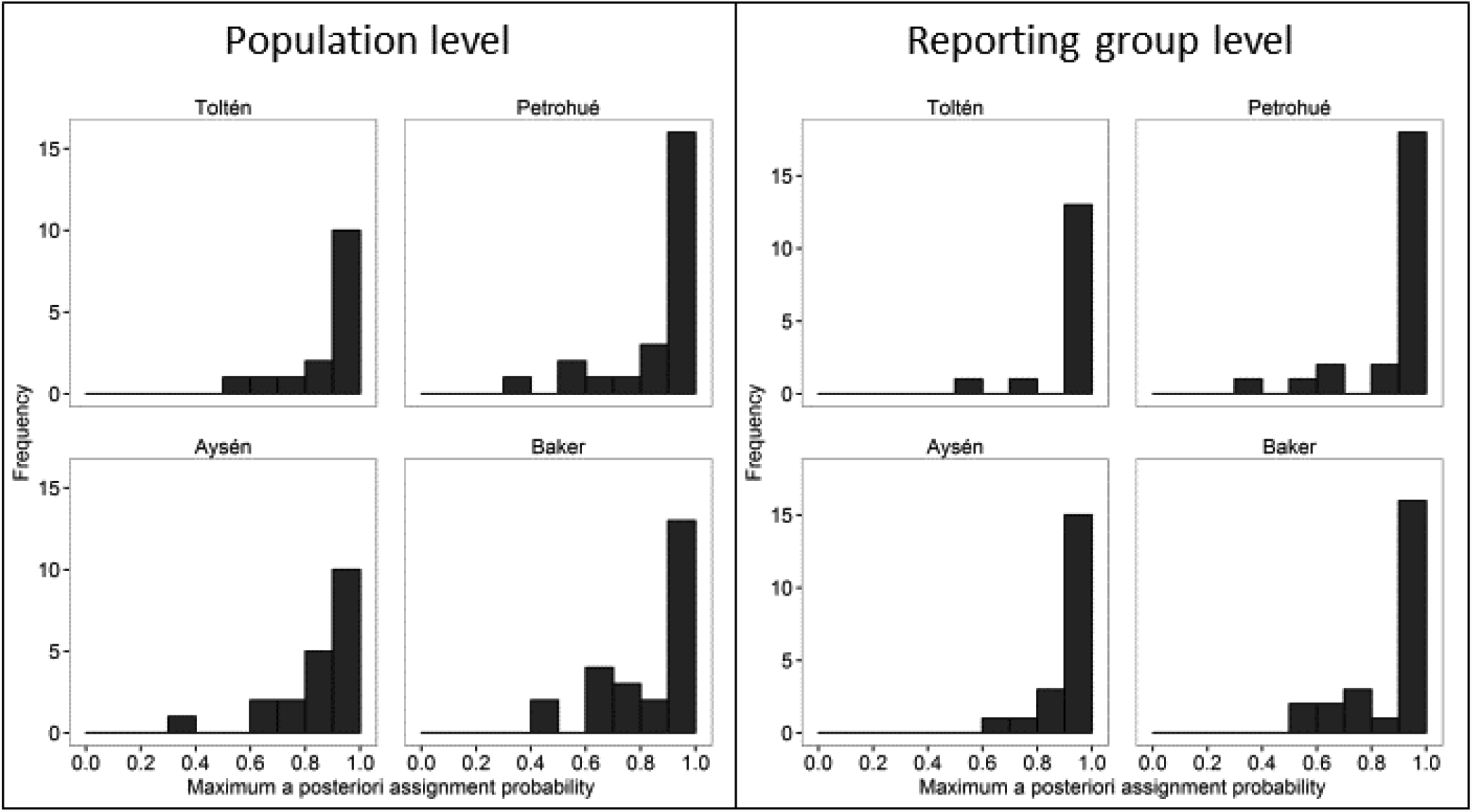
Frequency distribution of individual maximum assignment probabilities from population and reporting group-level CML mixture analysis.

**Figure S2:**
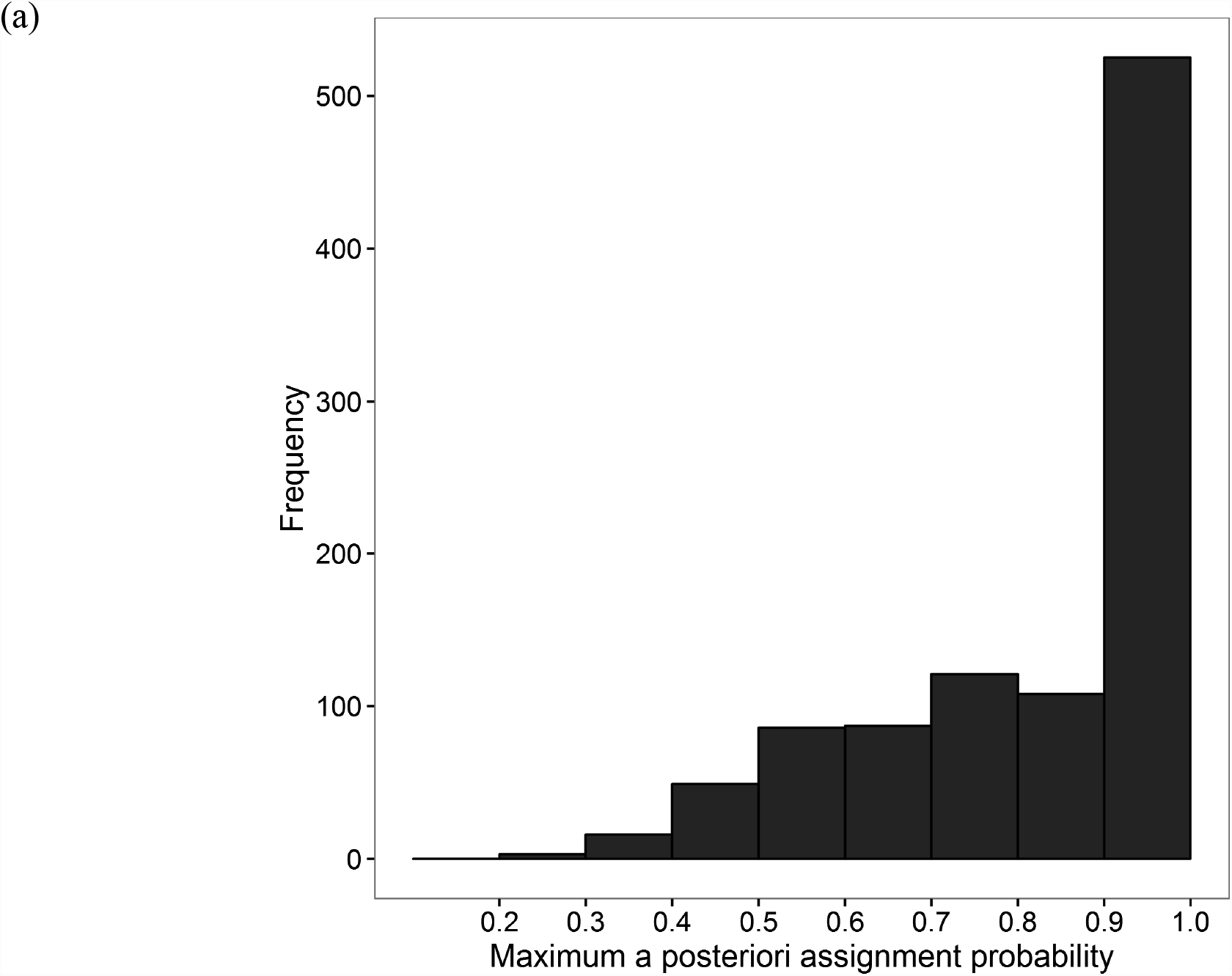
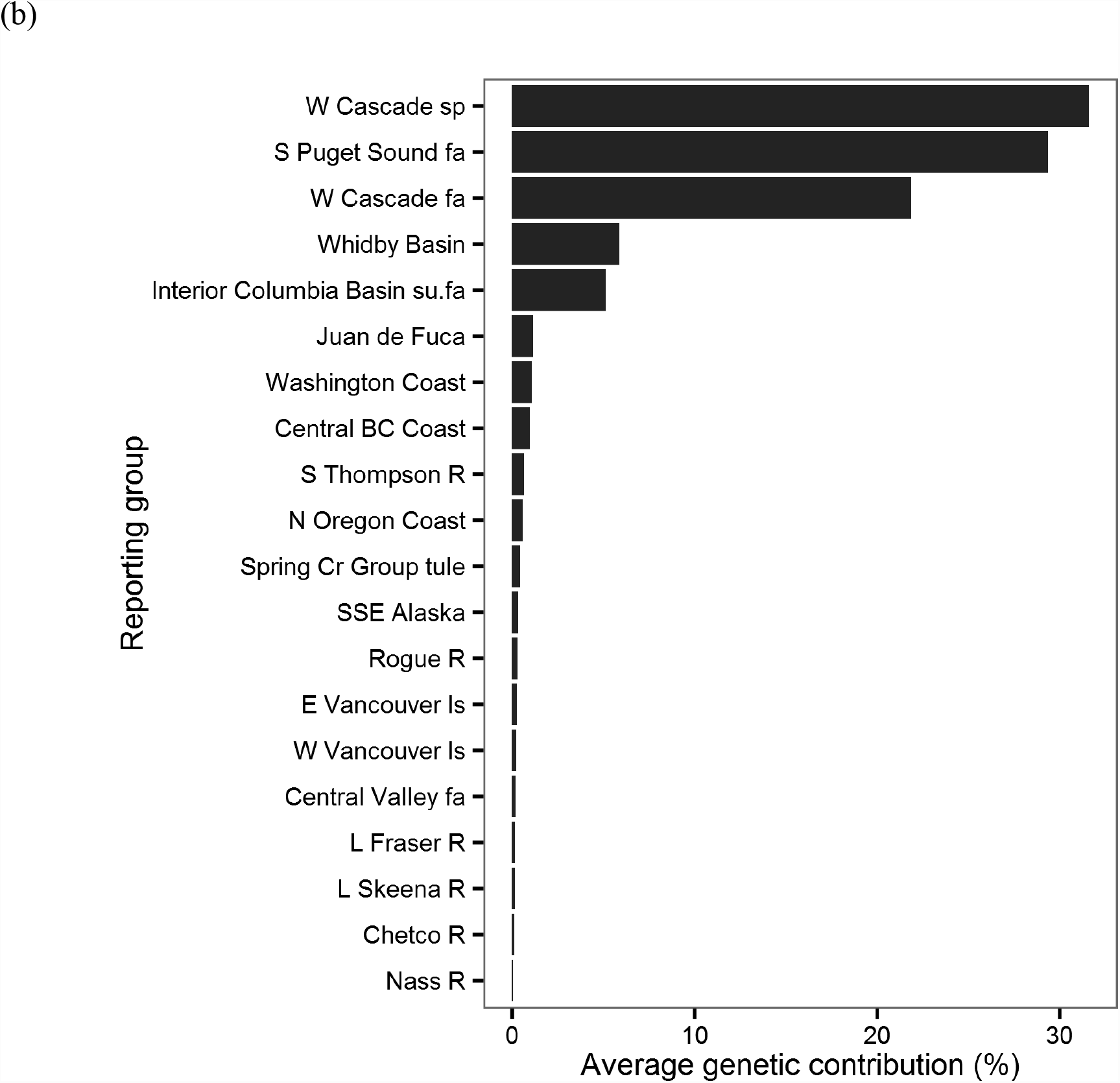
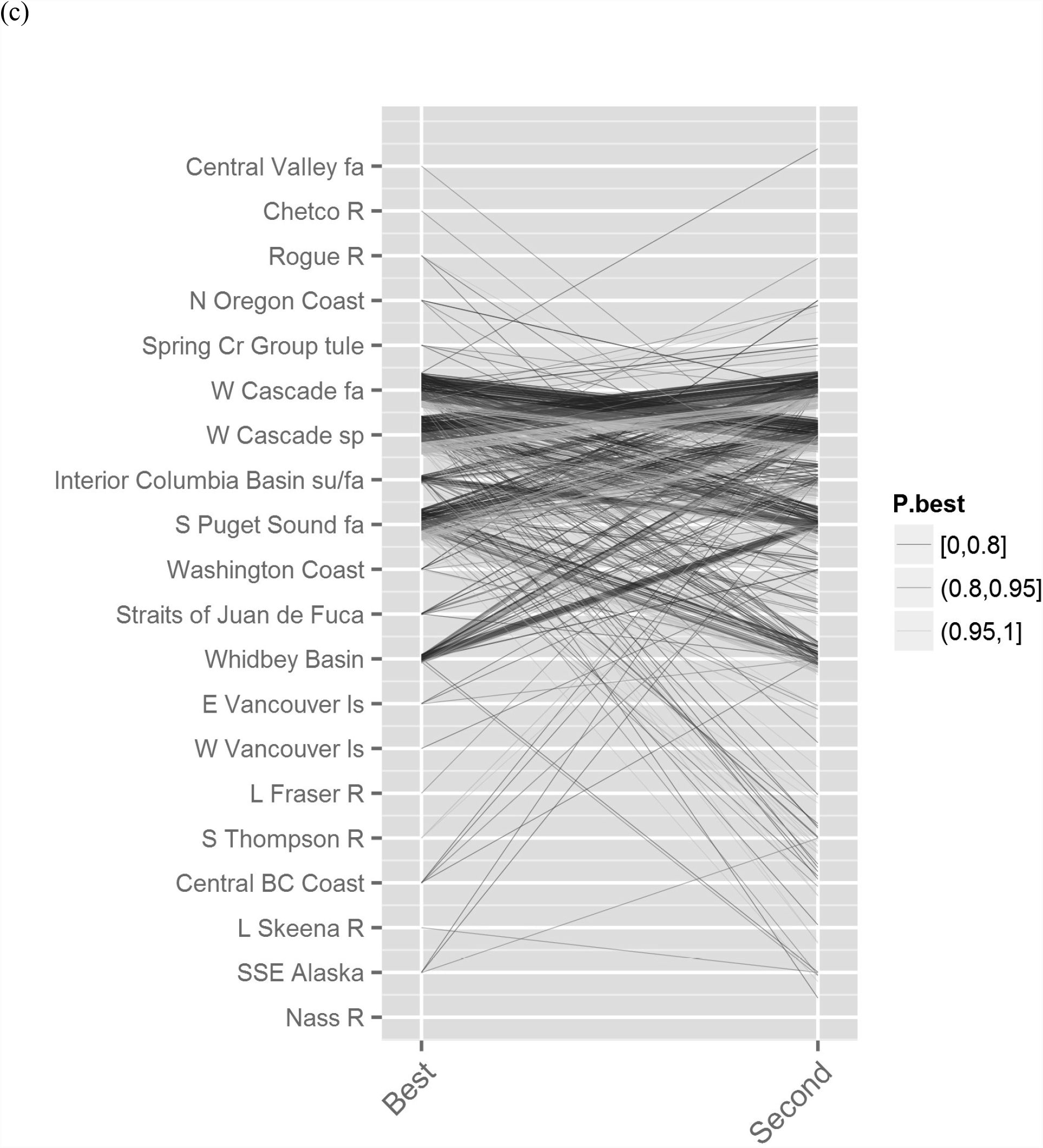
Conditional maximum likelihood (CML) mixture analysis of the simulated mixed-origin Chinook salmon population derived from Cowlitz River Hatchery spring run in the West Cascade spring-run reporting group and Soos Creek Hatchery fall run in the South Puget Sound fall-run reporting group. Distribution of maximum assignment probabilities (a), average percent genetic contribution of reporting groups to the simulated population (b), and individual alternative assignments based on best and second-best assignment probabilities (c). Equivocal assignments [i.e., low assignment probability, symbolized with darker lines in (c)] typically split probabilities between founder lineages, or between founder lineages and genetically similar reporting groups. A small fraction of simulated individuals assigned to unrelated lineages, even with high assignment probabilities in some cases. Reporting groups with no assignments were omitted. Reporting groups were ordered by decreasing order of estimated contribution (b) or increasing latitude (c).

**Figure S3:**
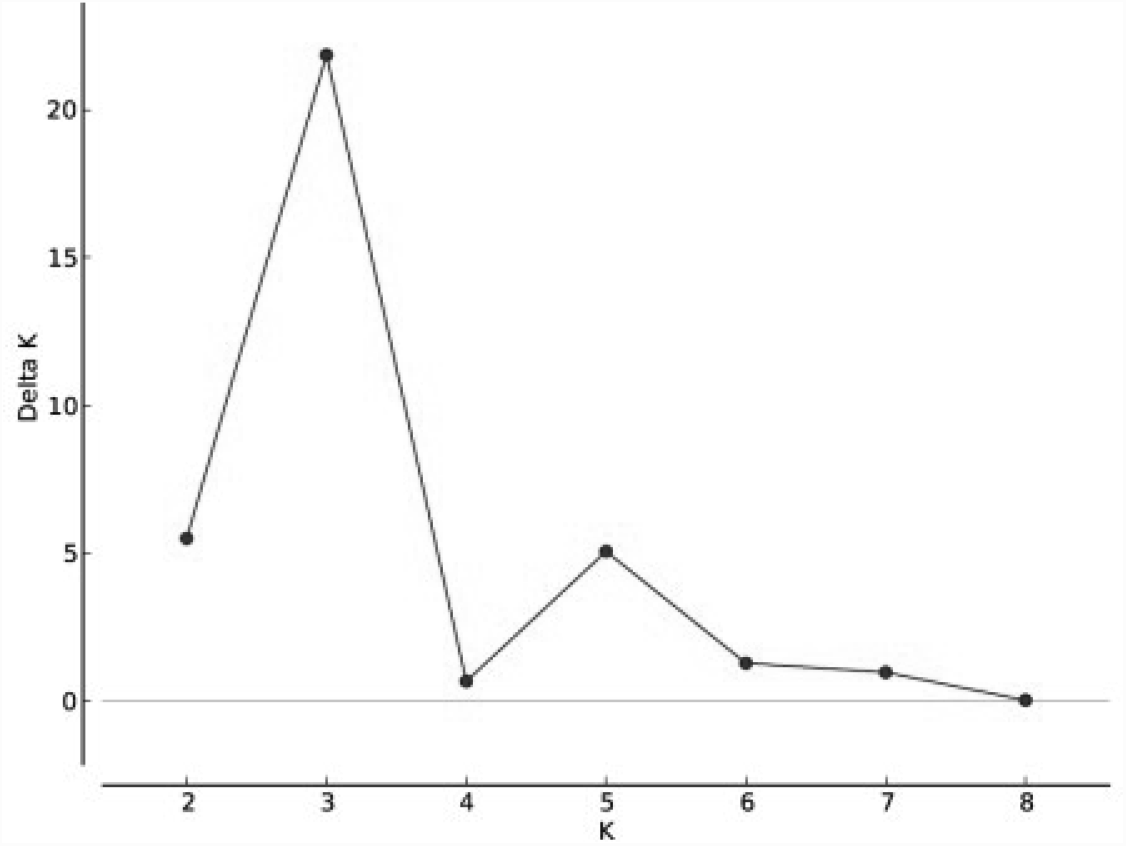
Delta K plot to determine appropriate number of groups in model-based clustering present in 8,228 fish from 31 populations distributed among seven North American lineages that potentially contributed founders of Patagonian populations.

**Figure S4:**
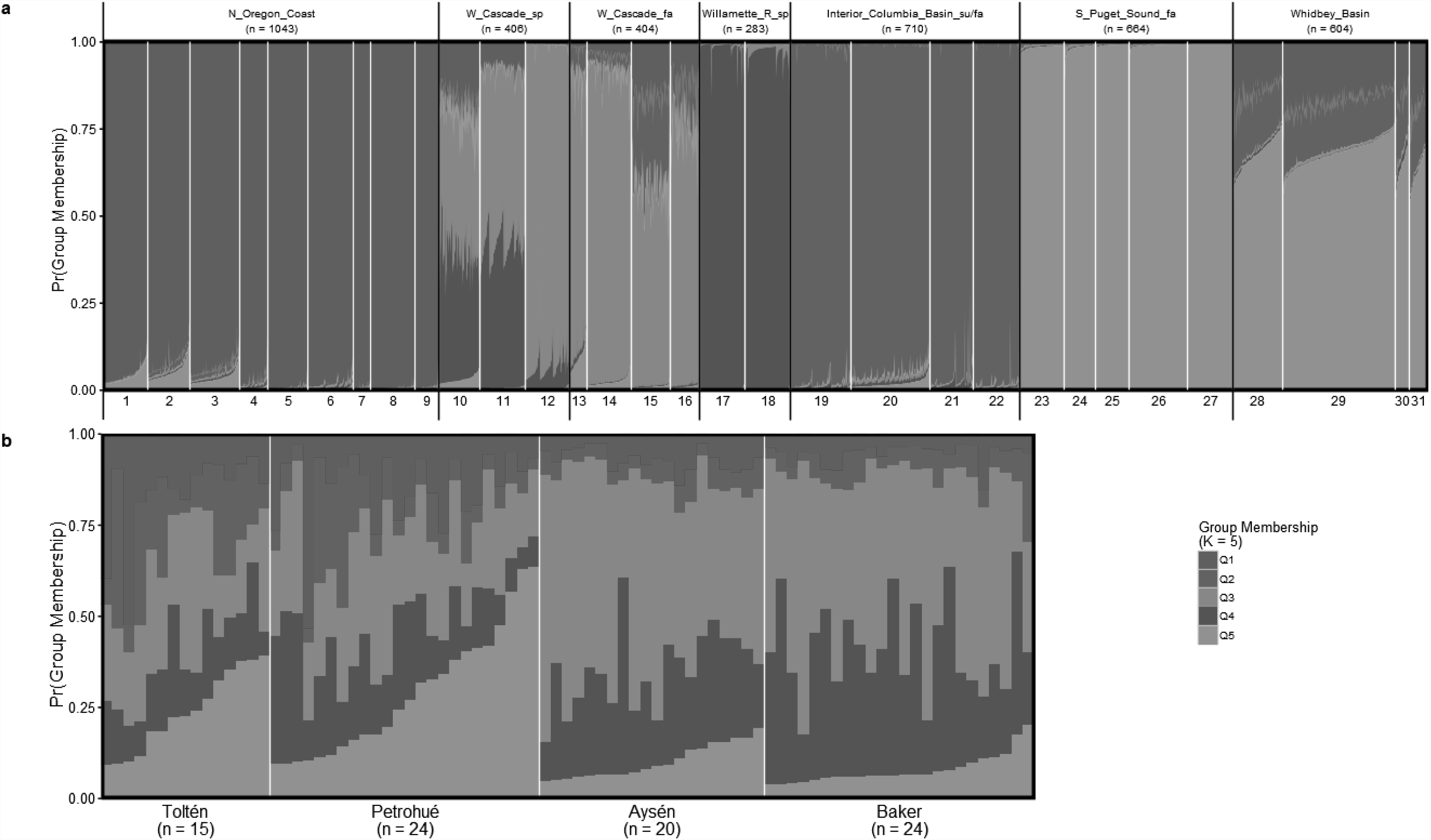
Individual stacked bar plot (STRUCTURE plot) of (a) North American baseline data-set of 8,228 fish from 31 populations distributed among seven North American lineages, and (b) Patagonian samples treated as having unknown origin in the analysis, and plotted by watershed (n = 81). The number of groups was set to *K* = 5, and estimated individual group membership probabilities is shown in colours.

